# Chikungunya virus live-cell reporter systems reveal cell-type-dependent antiviral responses to ribavirin

**DOI:** 10.64898/2026.10.27.762381

**Authors:** Samuel J. Dobson, Molly G. A. Brereton, Lai Wei, Christopher W. Wasson, Francesco Del Galdo, Andrew K. Tuplin

## Abstract

Chikungunya virus (CHIKV) remains a significant global public health threat, yet antiviral discovery is limited by experimental systems that provide little mechanistic resolution and rely heavily on transformed cell lines of uncertain physiological relevance. Here, we developed complementary live-cell CHIKV reporter systems comprising a dual-fluorescent subgenomic replicon and fluorescent infectious viruses that enable real-time monitoring of distinct stages of the viral replication cycle. By coupling an nsP3-tagged reporter to genomic RNA translation and replication with a subgenomic reporter of structural gene expression, we achieved temporal resolution of early and late replication events in living cells. Functional validation using replication-defective and subgenomic RNA-deficient mutants confirmed that the system distinguishes between viral RNA replication and downstream structural gene expression.

Application of these reporters revealed distinct antiviral responses across transformed and physiologically relevant human cell models. Ribavirin and the nsP2 inhibitor RA-0002034 potently inhibited CHIKV replication in transformed cell lines. Unexpectedly, ribavirin activity was markedly diminished in primary human skeletal muscle myoblasts and dermal fibroblasts, despite robust antiviral effects in commonly used screening cell lines. Mechanistic studies showed that ribavirin-mediated inhibition was rescued by guanosine but not inosine supplementation and was recapitulated by mycophenolic acid, consistent with guanine nucleotide depletion as the predominant antiviral mechanism under the conditions tested. Notably, inhibition through this pathway was substantially less effective in primary human cells.

Together, these findings establish versatile live-cell CHIKV reporter systems for mechanistic studies of viral replication and reveal striking cell-type-dependent differences in response to guanine nucleotide depletion. Our results demonstrate that host-cell context can profoundly influence both CHIKV replication and antiviral susceptibility, highlighting the importance of physiologically relevant models for understanding virus-host interactions and antiviral mechanism.

**Author Summary:** Chikungunya virus is a mosquito-borne pathogen that causes fever, severe joint pain, and chronic musculoskeletal disease. Although considerable progress has been made in understanding the viral replication cycle, important questions remain regarding how interactions between the virus and its host cell influence replication and susceptibility to antiviral intervention. Experimental systems that permit real-time analysis of distinct stages of infection are therefore valuable tools for advancing our understanding of chikungunya virus biology.

In this study, we developed complementary fluorescent chikungunya virus reporter systems that enable live-cell monitoring of viral replication and gene expression. These tools distinguished between early replication-associated processes and downstream subgenomic gene expression, providing mechanistic resolution of the viral replication cycle in living cells. Using these systems, we investigated the activity of ribavirin and found that its antiviral effects were highly dependent on host-cell context. While ribavirin potently inhibited chikungunya virus replication in transformed cell lines, this activity was largely absent in primary human muscle and skin cells that represent physiologically relevant targets of infection. Mechanistic studies showed that inhibition was linked to depletion of intracellular guanine nucleotides and could be reversed by guanosine supplementation. Together, our findings reveal how host-cell-dependent processes influence both chikungunya virus replication and antiviral susceptibility and provide new tools for dissecting virus-host interactions.

## 1. Introduction

Chikungunya virus (CHIKV) is an alphavirus of the *Togaviridae* family transmitted primarily by *Aedes* spp mosquitoes in tropical and subtropical regions (Weaver 2014; Weaver and Forrester 2015). Climate-change driven expansion of vector habitats has contributed to increasing global incidence and geographic spread of CHIKV transmission (Sharif et al. 2021; Ribeiro Dos Santos et al. 2025). The 2004-2006 Indian Ocean epidemic illustrates this expansion, with over 1.5 million cases reported (Sharif et al. 2021). More recent outbreaks, including the 2024–2025 epidemic on La Réunion Island (>51,000 confirmed and >188,600 cases) highlight the continuing public health burden (Frumence et al. 2025). Although mortality is low, up to 40-60% of infected individuals develop chronic disease characterised by persistent arthralgia and long-term morbidity (Schilte et al. 2013; Ng et al. 2024). In the absence of specific antiviral therapeutics, treatment remains largely supportive (Martelossi-Cebinelli et al. 2025). A live-attenuated CHIKV vaccine (IXCHIQ) was recently approved in the United States and European Union but safety concerns and reliance on surrogate efficacy endpoints have led to significant restriction or removal from use ((FDA) 2023, 2025a; (EMA) 2024; (UKHSA) 2026). A recombinant virus-like particle vaccine (VIMKUNYA) has more recently been approved; however, the continued absence of approved antiviral therapies highlights the ongoing need for effective antiviral strategies ((EMA) 2025; (FDA) 2025b). High-content screening (HCS) platforms therefore represent an important approach for identifying candidate antivirals and expanding the currently limited CHIKV therapeutic pipeline, particularly given the absence of clinically approved small molecule inhibitors (SMIs) (de Souza et al. 2024). However, an additional challenge is determining whether antiviral activities identified in conventional transformed-cell screening models are maintained in physiologically relevant human target cells.

CHIKV has an ∼11.8 kb 5¢ type-O-N-methylguanosine capped and 3¢ poly(A) tailed positive-sense single-stranded RNA genome, that is encapsidated by an enveloped virion (Kendall et al. 2019). The genome consists of two open-reading frames (ORFs) that are flanked by highly structured untranslated regions which regulate replication efficiency (Sun et al. 2024; Kendall et al. 2019; Remenyi et al. 2018; Prosser, Stonehouse, and Tuplin 2023). ORF-1 encodes the non-structural proteins (nsPs) 1-4 that form the viral replication machinery, whilst ORF-2 encodes the structural proteins capsid, E1, E2, E3 and 6K/TF (Gao et al. 2019). Translation of ORF-1 initiates directly from genomic RNA (gRNA), whereas ORF-2 is translated from a sub-genomic RNA (sgRNA) that is transcribed from a negative-strand intermediate produced during CHIKV genome replication by the viral polymerase nsP4 (Kendall et al. 2019; Muller et al. 2019).

Despite the availability of replication-competent CHIKV infectious clones encoding fluorescent tags, recent developments in cell-based antiviral screening of CHIKV have relied on immunofluorescence assays (IFA) (Sawetpiyakul et al. 2025; Colunga-Saucedo et al. 2022; Tsetsarkin et al. 2006). However, antibody-based detection introduces multiple processing steps, batch variability and increased cost, limiting scalability for HCS. Furthermore, most CHIKV HCS approaches are constrained by single-endpoint measurements that collapse the viral life cycle into a single readout, limiting the ability to distinguish inhibitors of entry, genome replication and sgRNA translation. The use of sub-genomic replicons (SGRs) offers the ability to study CHIKV genome replication and translation independent of entry and virion formation, supporting mechanistic investigation in biosafety level 2 (BSL-2) laboratories and permitting greater accessibility for screening. To date, CHIKV SGR systems have mostly relied on luciferase assays, which whilst providing good sensitivity do not allow real-time monitoring of replicase activity, spatial dynamics of replication complexes or temporal resolution of antiviral effects (Pohjala et al. 2011; Lello et al. 2020; Yin et al. 2026). The requirement for fixed time-point lysis sampling and downstream processing limits the HCS potential of such assays, while the resulting lack of kinetic resolution obscures early replication events, complicates assignment of compound mechanism-of-action and prevents mechanism-aware triaging of antiviral hits, thereby hindering CHIKV drug discovery.

Fluorescent imaging platforms have proven transformative for studying viral replication across a wide range of viruses including enteroviruses, hepatitis C virus, polyomaviruses and lymphocytic choriomeningitis arenavirus, however many of these systems have not fully exploited the potential for real-time dissection of replication dynamics (Dobson, Mankouri, and Whitehouse 2020; Dobson et al. 2023; Kejriwal et al. 2023; Ward et al. 2024; Stewart et al. 2015; Byford et al. 2024). Previous approaches to study CHIKV have largely relied on single reporters or static imaging, which do not enable simultaneous monitoring of multiple stages of infection. Although fluorescent reporter systems have been developed for alphaviruses such as Sindbis virus, no platform currently enables simultaneous, real-time monitoring of both genomic and sub-genomic CHIKV gene expression (Jose, Taylor, and Kuhn 2017). Here, we report the development of a spectrally distinct, dual-fluorescent, CHIKV SGR system incorporating an nsP3-mNeonGreen (nsP3-mNG) fusion and mScarlet-I3 replacement of ORF-2, which coupled with IncuCyte live-cell imaging this system permits real-time multiplexed monitoring of viral replication dynamics. This system overcomes the temporal, throughput and antibody-dependent limitations of existing SGR- and IFA-based platforms by enabling continuous, multiplexed quantification of CHIKV replication dynamics in living cells. Although prior studies have characterised replication-complex architecture and nsP3-linked granule dynamics using static or single-reporter systems, no published assay provides dual-colour, real-time visualisation of both ORF-1 and ORF-2 translation (Kendall et al. 2019; Remenyi et al. 2018; Gao et al. 2019; Kril et al. 2024). This platform therefore establishes a framework for mechanistic investigation of CHIKV replication dynamics and antiviral activity. Importantly, despite decades of investigation, the mechanisms underlying the antiviral activity of many compounds evaluated against CHIKV remain incompletely understood. Ribavirin (RIBA), one of the most widely used broad-spectrum antivirals tested against alphaviruses, has been reported to inhibit CHIKV replication in multiple experimental systems, yet the dominant mechanism responsible for this activity remains unresolved. Proposed models include inhibition of inosine monophosphate dehydrogenase (IMPDH) and depletion of intracellular guanine nucleotide pools, inhibition of equilibrative nucleotide transporter 1 (ENT1), direct effects on viral RNA synthesis and mutagenic error catastrophe resulting from incorporation into nascent viral RNA (Franco et al. 2018; Shah, Sunderland, and Grdzelishvili 2010). The inability of conventional endpoint assays to distinguish between these possibilities highlights the need for platforms capable of linking antiviral efficacy with mechanistic insight during infection and across different cellular contexts.

In addition, we extend this platform through the generation of spectrally distinct infectious CHIKV clones expressing nsP3-mNG or nsP3-mScarlet-I3 (nsP3-mS3). Whilst the SGR enables multiplexed, real-time resolution of ORF-1 and ORF-2 genome replication and translation dynamics under BSL-2 containment, the infectious clone systems permit full life-cycle analysis using a choice of fluorophores that can be matched to assay requirements or used to circumvent compound-dependent autofluorescence and spectral interference during screening (Pohjala et al. 2011). Together, findings from the current study demonstrate that the platform overcomes the limitations of fixed endpoint assays by enabling real-time, mechanism-aware interrogation of CHIKV replication dynamics. Beyond assay development, application of the system enables direct comparison of antiviral efficacy across conventional screening models and physiologically relevant human target cells, while providing mechanistic insight into compound activity. In addition to revealing distinct antiviral and replication phenotypes across transformed, immortalised and primary human cell models, including the reduced efficacy of RIBA in physiologically relevant human skeletal muscle myoblast (HSkMM), primary human dermal fibroblast (pHDF) and hTERT-immortalised human dermal fibroblast (hTERT-HDF) systems, the platform also enables direct interrogation of antiviral mechanism. By resolving the effect of antiviral intervention on both ORF-1- and ORF-2-associated processes, the system provides a means to link phenotypic inhibition with specific stages of CHIKV replication.

Collectively, the dual-colour SGR and infectious reporter viruses establish a complementary live-cell imaging platform that supports antiviral discovery while simultaneously providing mechanistic insight into both compound activity and the cell-type-dependent determinants of antiviral efficacy.

## 2. Results

### 2.1. Development of a live-cell CHIKV dual-fluorescent sub-genomic reporter assay

To enable dynamic, multiplexed monitoring of CHIKV replication, we developed a fluorescent SGR system in which mNG was inserted into a permissive region of the nsP3 hypervariable domain, previously shown to tolerate reporter fusion without impairing replicase activity (Pohjala et al. 2011). In parallel, ORF-2 was replaced with mS3, enabling simultaneous detection of genomic and sub-genomic RNA translation (Figure 1A). mNG and mS3 were selected because of their higher quantum yield, enhanced photostability, rapid maturation and improved spectral separation for live-cell imaging in comparison to alternative fluorescent proteins.

**Figure 1:**
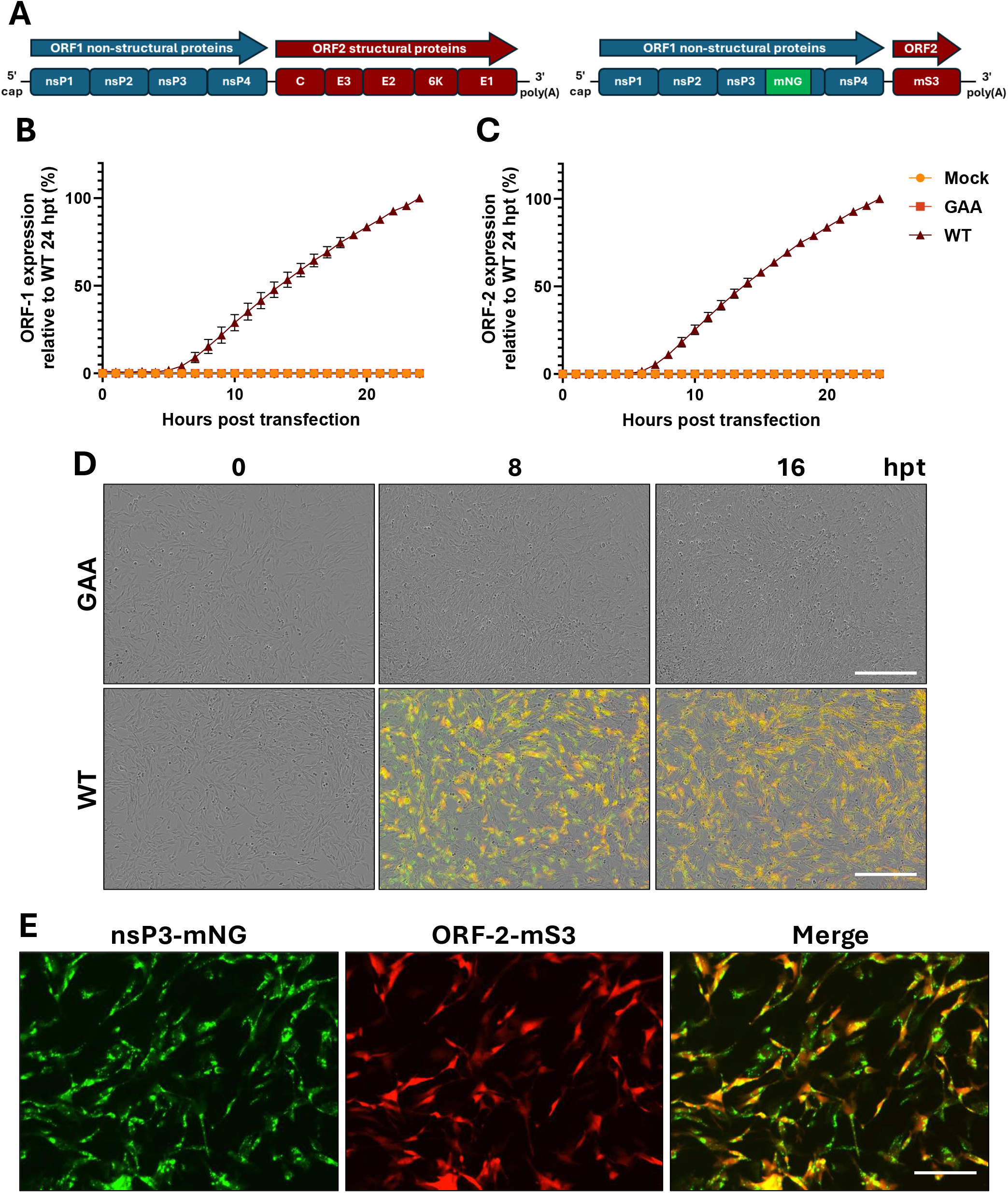
Characterisation of the dual-fluorescent CHIKV sub-genomic replicon. (A) schematic representation of WT genomic and sub-genomic replicon architecture, indicating insertion of mNG within nsP3 (ORF-1) and replacement of the ORF-2 coding region with mS3. (B-C) Live-cell, hourly quantification of nsP3-mNG (ORF-1) and ORF-2-mS3 expression following transfection of BHK-21 cells with WT or replication-defective (GAA) sub-genomic replicon RNA. Fluorescence intensity was normalised to 24 hpt. (D) Representative images of BHK-21 cells transfected with WT or GAA sub-genomic replicon RNA at 0, 8 and 16 hpt. Scale bars 400 µm. (E) Magnified image of BHK-21 cells at 8 hpt following transfection with WT sub-genomic replicon RNA, highlighting nsP3-mNG puncta. Scale bar 100 µm. Data are presented as mean ± SEM (n = 3 independent experiments)

A replication-defective control replicon was generated by substituting the nsP4 catalytic GDD motif with GAA, a well-established polymerase-inactivating mutation used in CHIKV and other alphavirus replicons to distinguish input translation from bona fide RNA replication (Pohjala et al. 2011).

Following transfection of BHK-21 cells, translation reached a detectable level 4h post transfection (hpt), with robust translation observed from 5 hpt for wild-type (WT) replicons, whilst GAA controls showed no detectable fluorescence (Figure 1B-D). Fluorescent signal for both nsP3-mNG and ORF-2-mS3 continued to increase progressively over time, with WT trajectories deviating from linear progression approximately 16 hpt for both ORF-1 (Figure 1B) and ORF-2 (Figure 1C).

During live-cell imaging, nsP3-mNG formed characteristic punctate cytoplasmic foci (Figure 1E), consistent with nsP3-containing replication granules previously shown to represent tubular helical assemblies essential for alphavirus RNA synthesis (Kril et al. 2024). In contrast, ORF-2-mS3 remained diffuse throughout the cell, supporting its utility as a reporter of sgRNA-driven translation. Together, these data confirmed that the dual-fluorescent SGR enabled real-time monitoring of genomic and sub-genomic translation while simultaneously resolving replication-complex dynamics in living cells.

### 2.2. nsP3 PV247/8AA mutation confirms decoupling of ORF-1 and ORF-2 translation

To validate the ability of the SGR system to resolve distinct stages of the CHIKV replication cycle, reporter dynamics were examined using a construct containing the nsP3 mutation PV247/8AA. This mutation has previously been shown to impair sgRNA production without abolishing viral gRNA replication, providing an established benchmark for functional separation of gRNA and sgRNA synthesis (Gao et al. 2019). Based on the kinetics of CHIKV SGRs established earlier, discrete early (8 hpt) and late (16 hpt) time points were selected to assess replication complex formation corresponding to non-structural protein expression, and robust sgRNA transcription and translation of ORF-2 encoded products, respectively.

Consistent with previous reports, PV247/8AA replicons retained substantial nsP3-mNG signal following transfection, indicating that input RNA translation and replication-associated non-structural protein expression were largely preserved (Figure 2A). Although the kinetics of nsP3-mNG accumulation were modestly delayed relative to WT, fluorescence continued to rise throughout the time course and reached 77% of WT by late time points (Figure 2C). These data indicate that nsP3-mNG accurately reports replicase-associated activity while remaining sensitive to changes in replication efficiency, due to replication-defective mutations.

**Figure 2:**
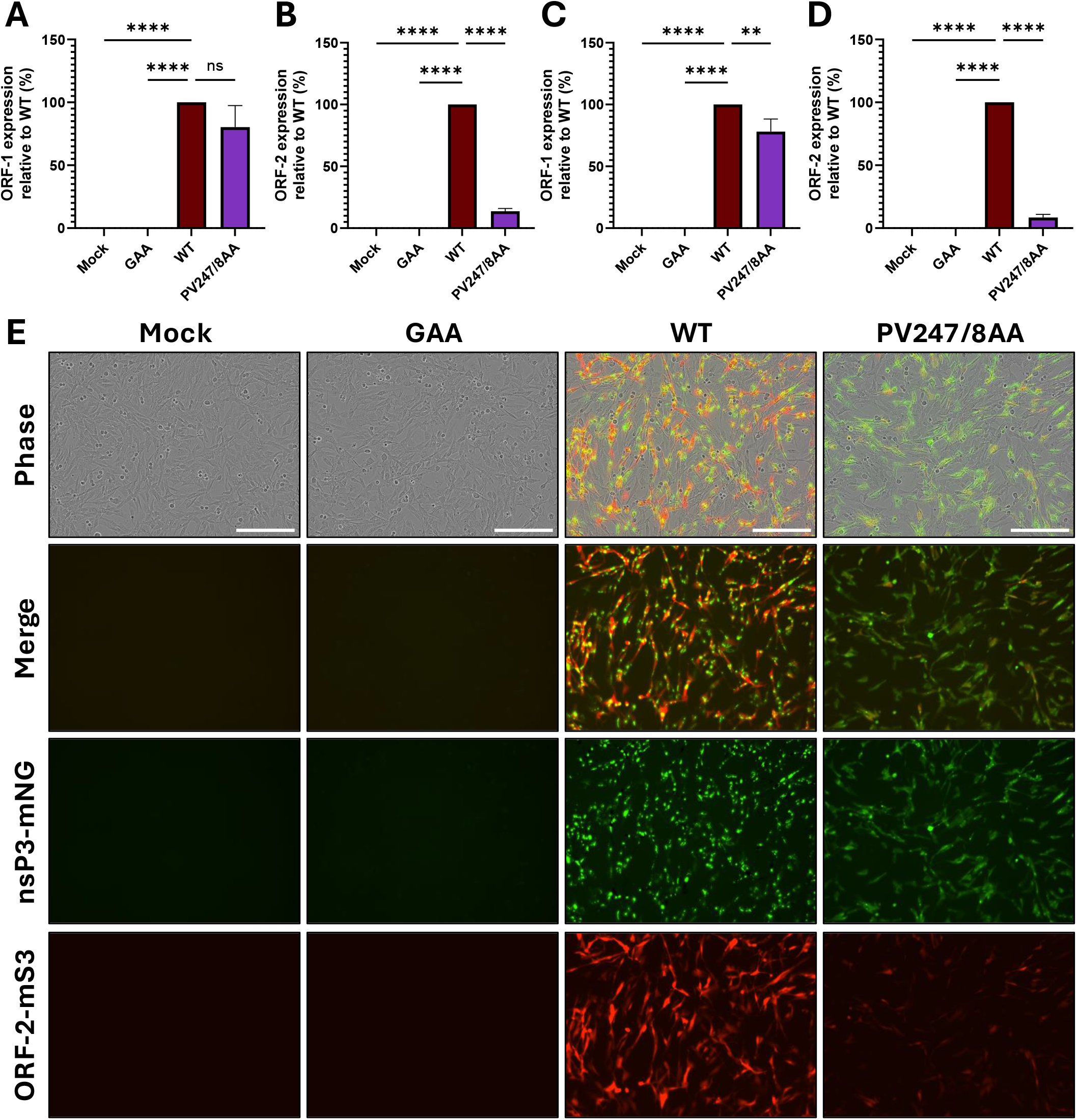
Validation of replication stage separation using the nsP3 PV247/8AA mutant. (A-D) BHK-21 cells were transfected with GAA, WT or PV247/8AA CHIKV sub-genomic replicon RNA and imaged at 8 and 16 hpt. Expression of ORF-1 (nsP3-mNG) is shown in (A, C), and expression of ORF-2-mS3 is shown in (B, D) at 8 and 16 hpt, respectively. (E) Representative images of BHK-21 cells transfected with GAA, WT or PV247/8AA sub-genomic replicon RNA at 16 hpt. Scale bar 200 µm. Data are presented as mean ± SEM (n = 3 independent experiments).

In contrast, expression of ORF-2-mS3 was profoundly impaired in the PV247/8AA replicon (Figure 2B, D). Whereas WT replicons showed robust accumulation of mS3 fluorescence following an initial lag phase, PV247/8AA showed minimal red fluorescence throughout the assay, reaching only 14% of WT signal at endpoint. Importantly, this defect was not consistent with a simple temporal delay, as mS3 fluorescence remained strongly suppressed, even at late time points where nsP3-mNG was readily detectable. These findings demonstrate that the ORF-2-mS3 reporter selectively reports sgRNA-driven translation and is highly sensitive to perturbations in sgRNA synthesis.

Consistent with previous reports, PV247/8AA nsP3-mNG remained diffuse throughout the cytoplasm rather than forming puncta, as observed for WT nsP3-mNG (Figure 2E). This divergence provides direct evidence that the dual-fluorescent SGR not only discriminates between early replication-associated events and later sgRNA-dependent translation but also provides information regarding nsP3 puncta formation.

### 2.3. Validation of the CHIKV SGR platform for HCS of small-molecule inhibitors

To assess the suitability of the dual-fluorescent SGR for HCS, we evaluated the antiviral activity of the well-characterised inhibitor RIBA and the nsP2-targeting RA-0002034 (RA). RIBA is a broad-spectrum antiviral nucleoside analogue with demonstrated inhibitory activity against numerous RNA viruses, including CHIKV. Although the exact mechanism of CHIKV inhibition has not been identified, it is suggested to occur through inhibition of ENT1, IMPDH inhibition, RNA mutagenesis and/or interference with viral RNA synthesis (Tam, Lau, and Hong 2001; Bougie and Bisaillon 2004). Consistent with previous reports of CHIKV inhibition in cell culture, RIBA displayed strong antiviral effects across the SGR reporters (Figure 3) (Gallegos et al. 2016). At 8 hpt, ORF-1 translation was inhibited by 75%, 96% and 100% following treatment with 5, 10 and 20 µM RIBA, respectively (Figure 3A), while ORF-2 translation was completely abolished at ≥10 µM and reduced by 95% at 5 µM (Figure 3B). At 16 hpt, a clear dose-dependent reduction in reporter accumulation was observed for both ORF-1 and ORF-2 (Figure 3CCD), with inhibition of 34–80% for ORF-1 and 48–94% for ORF-2.

**Figure 3:**
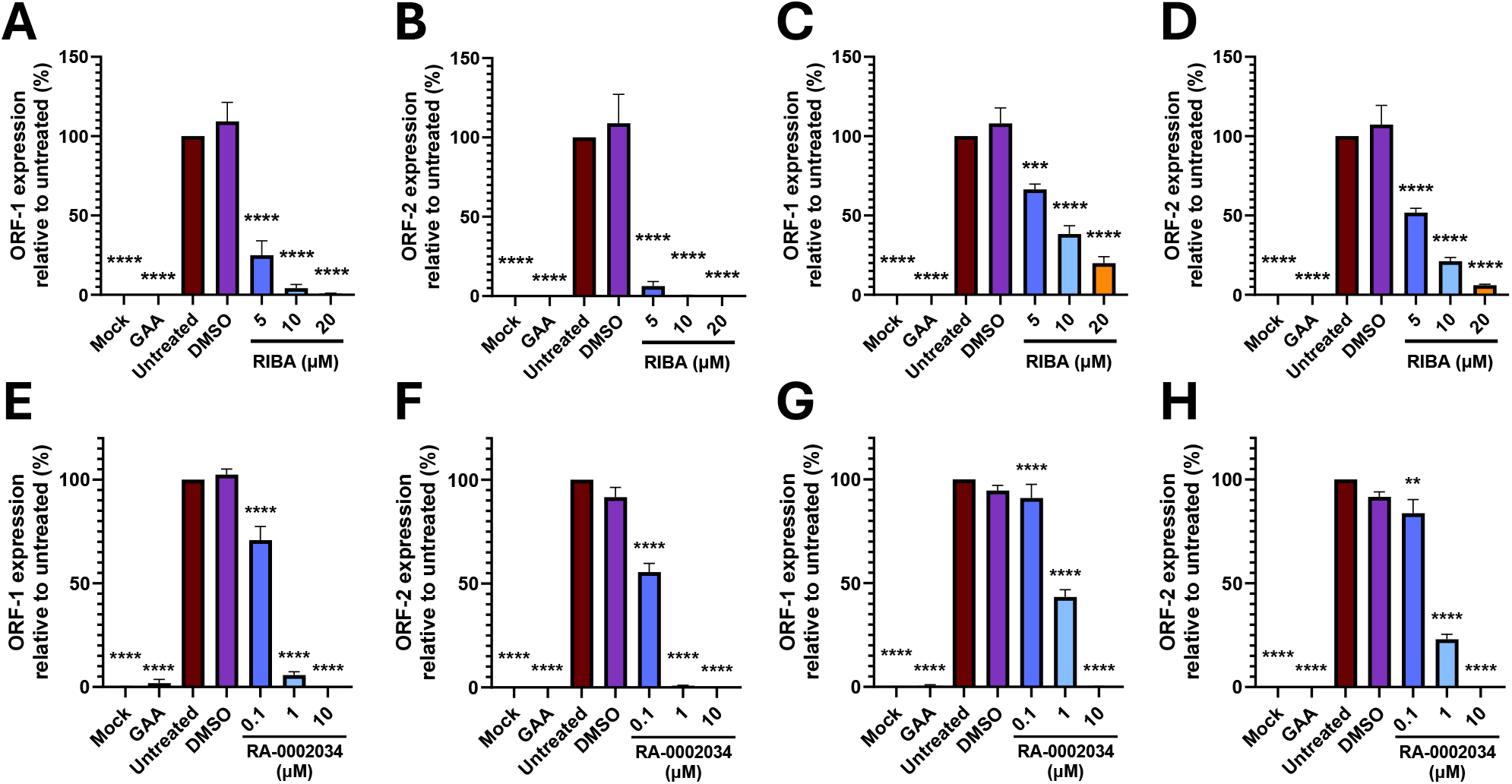
Validation of the CHIKV sub-genomic replicon system for high-throughput screening. BHK-21 cells were transfected with WT CHIKV sub-genomic replicon RNA in the presence of RIBA at 5, 10 and 20 µM (A-D) or RA-0002034 at 0.1, 1 and 10 µM (E-H). DMSO was included as a vehicle control, alongside untreated, GAA and non-transfected (mock) controls. ORF-1 (nsP3-mNG) and ORF-2-mS3 expression were analysed at 8 hpt (A, B, E, F) and 16 hpt (C, D, G, H). Data are presented as mean ± SEM (n = 3 independent experiments).

RA treatment similarly resulted in a clear, dose-dependent suppression of both ORF-1 and ORF-2 (Figure 3E-H). RA covalently modifies the catalytic cysteine of the nsP2 cysteine protease domain and does not show significant off-target inhibition (Merten et al. 2024; Hossain et al. 2025). At 8 hpt, inhibition was partial at 0.1 µM (29% reduction, ORF-1; 44% reduction, ORF-2), near-complete at 1 µM and complete at 10 µM for both reporters. At 16 hpt, ORF-1 expression had recovered to WT levels in the presence of 0.1 µM RA, whilst ORF-2 was only 17% reduced. Partial recovery was observed with 1 µM, with 57% inhibition for ORF-1 and ORF-2, respectively. Treatment with 10 µM maintained complete suppression of replication across ORF-1 and ORF-2. Notably, the reduction of ORF-2 signal paralleled that observed for ORF-1, confirming that RA impacts upstream replication processes rather than impairing sgRNA synthesis and translation. This similarity in inhibition profiles between the two reporters is consistent with targeting of nsP2-dependent functions required for replicase activity.

Together, these data demonstrate that the dual-fluorescent SGR reliably recapitulates known antiviral phenotypes, while providing temporal resolution unavailable in conventional endpoint luciferase or IFAs. By allowing parallel, real-time quantification of genomic and sub-genomic translation, the system enables mechanistic evaluation of CHIKV inhibitors and prioritisation of antiviral candidates.

### 2.4 Characterisation of fluorescently-tagged nsP3 infectious clones

Existing CHIKV infectious clone systems incorporating fluorescent reporters often utilise fluorophores with suboptimal brightness and maturation kinetics, limiting sensitivity for real-time imaging of replication dynamics. To improve signal quality, mNG and mS3 were inserted in parallel into the previously described permissive site within the nsP3 hypervariable domain (Figure 4A) (Pohjala et al. 2011; Remenyi et al. 2017).

**Figure 4:**
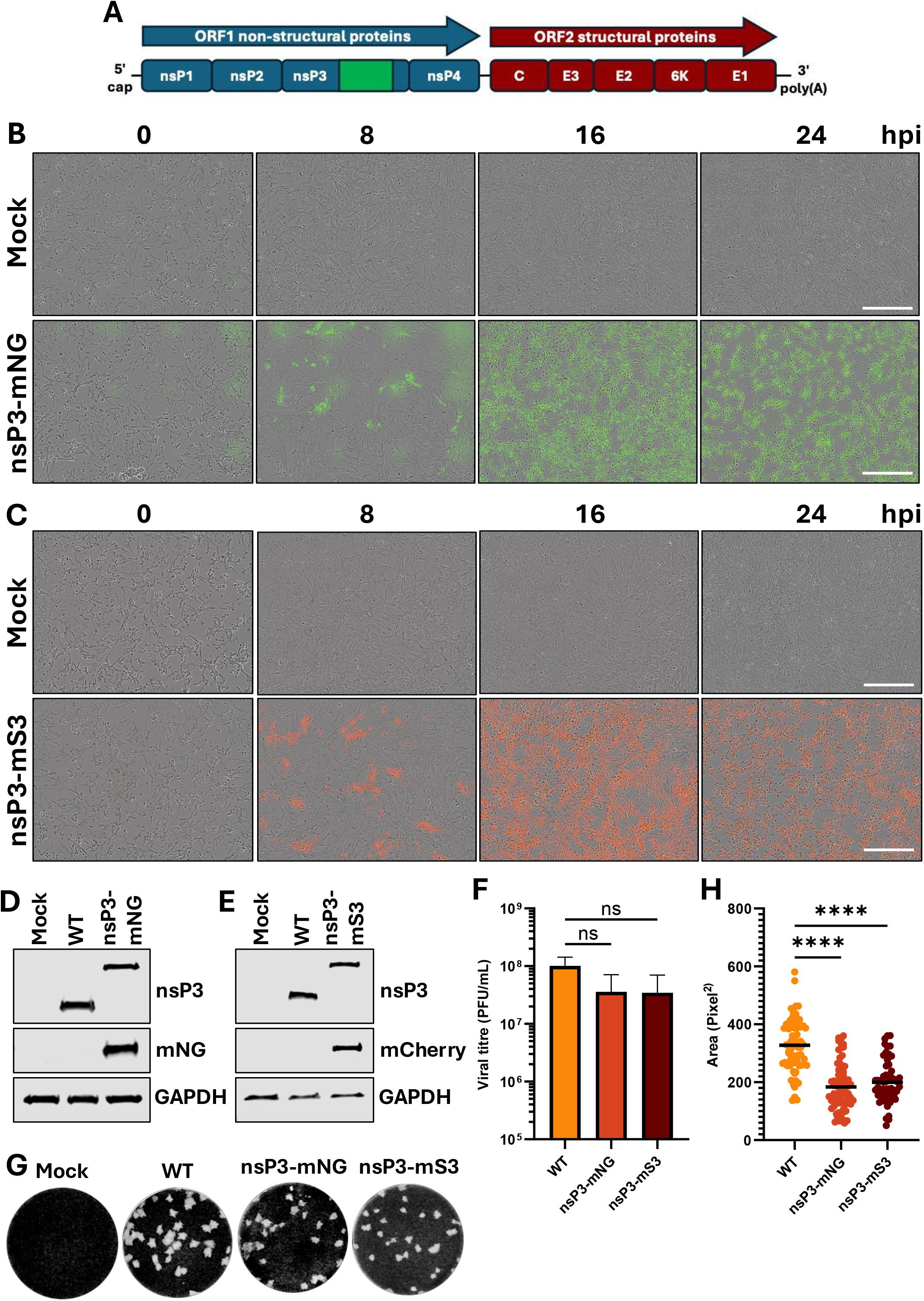
Development and validation of nsP3-tagged infectious clones. (A) Schematic representation of the CHIKV infectious clone architecture, indicating the insertion site of mNG or mS3 within nsP3. (B, C) Live-cell imaging of BHK-21 cells infected with nsP3-mNG or nsP3-mS3 virus at an MOI of 0.1, showing fluorescence at 0, 8, 16 and 24 hpi. Scale bars 400 µm. (D, E) Western blot analysis of nsP3 expression in BHK-21 cells at 16 hpi with WT, nsP3-mNG or nsP3-mS3 viruses. (F) Viral titres in supernatants collected from infected BHK-21 cells 16 hpi, determined by plaque assay. Data are presented as mean ± SEM. (G) Representative plaques formed by WT, nsP3-mNG and nsP3-mS3 viruses in BHK-21 cells at 72 hpi (dilutions shown: WT, 10-7; nsP3-mNG, 10-6; nsP3-mS3, 10-6). (H) Quantification of plaque size for WT, nsP3-mNG and nsP3-mS3 viruses. n = 3 independent experiments.

Following rescue of infectious virus, infection of BHK-21 cells confirmed that both nsP3-mNG and nsP3-mS3 constructs retained replication competence. Live-cell imaging revealed initial infection events at 8 h post-infection (hpi), followed by progressive direct cell-to-cell spread and widespread secondary infection from newly produced infectious virus at 16 and 24 hpi (Figure 4B, C). Replication kinetics were comparable between both reporter viruses, indicating that incorporation of either fluorophore did not substantially impair virus replication or spread.

Western blot analysis confirmed expression of the tagged nsP3 proteins with the expected ∼27 kDa mobility shift relative to WT nsP3, with no detectable reversion to the untagged form (Figure 4D, E). Immunoblotting using anti-fluorophore antibodies further verified the presence of mNG and mS3 (detected using an mCherry-cross-reactive antibody), confirming stable reporter incorporation. Stability of reporter incorporation was also tested by blind passage, with signal for mNG and mS3 remaining comparable across 10 passages (Supp. Figure 1A). Live-cell imaging further confirmed complete monolayer infection of reporter viruses 16 hpi (Supp. Figure 1B).

To assess the impact of reporter insertion on viral fitness, supernatants collected from low multiplicity (MOI 0.1) infections at 16 hpi were analysed by plaque assay. Both nsP3-mNG and nsP3-mS3 viruses exhibited a modest reduction in viral titre relative to WT; although, these differences were not statistically significant (Figure 4F). Consistent with this, plaque morphology analysis revealed a subtle reduction in plaque size for both tagged viruses compared with wild type (Figure 4G, H). Together, these data demonstrate that fluorescent tagging of nsP3 permits robust live-cell imaging of CHIKV infection while maintaining near-wild-type replication competence.

### 2.5. Antiviral screening using nsP3-mNG infectious CHIKV across multiple cell types

To evaluate the utility of the nsP3-tagged infectious clone platform for antiviral screening and the conservation of antiviral response across cell types, we assessed RIBA activity across three immortalised cell lines commonly used for CHIKV replication studies: BHK-21, HEK293T and Huh7 cells. Cells were infected with nsP3-mNG CHIKV at an MOI of 0.1 (BHK-21 and HEK293T) or 10 (Huh7), reflecting the lower permissivity of Huh7 cells, and treated with RIBA at 5–20 µM. Viral replication was quantified by mNG fluorescence, while confluence was simultaneously measured as an indicator of compound-associated cytotoxicity (Figure 5).

**Figure 5:**
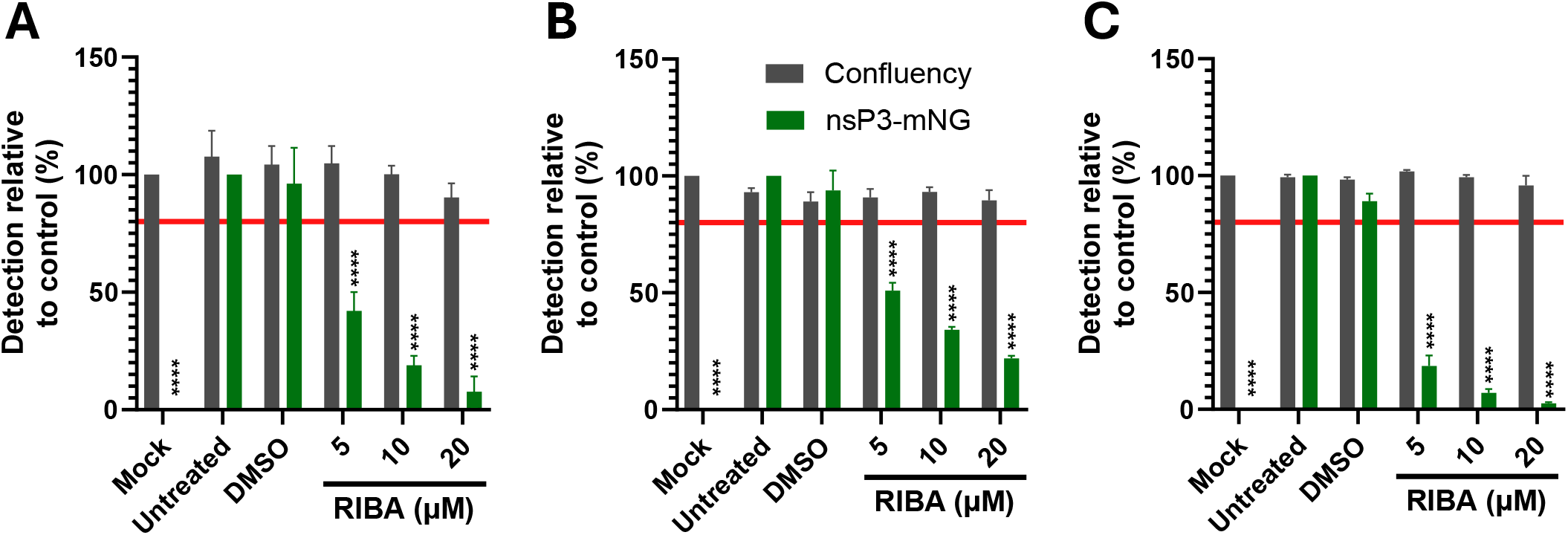
Screening of ribavirin antiviral activity against the CHIKV nsP3-mNG infectious clone in immortalised cell lines. BHK-21 (A), HEK293T (B), and Huh7 (C) cells were infected with CHIKV nsP3-mNG at a MOI of 0.1 (BHK and HEK293T) or 10 (Huh7), in the presence of RIBA at 5, 10, and 20 µM. Cells were incubated for 18 h (BHK) or 24 h (HEK293T and Huh7) prior to live-cell imaging. Viral replication was quantified by mNG fluorescence intensity, and cell confluency was measured in parallel as an indicator of compound-associated cytotoxicity. Red line denotes the 80% confluency threshold for acceptable cell viability. Black asterisks indicate statistically significant differences in viral replication relative to untreated infected controls. No significant difference in confluency was observed between mock and treated samples. Data are presented as mean ± SEM (n = 3 independent experiments).

In BHK-21 cells (Figure 5A), RIBA produced a strong, dose-dependent reduction in viral replication with minimal effects on confluence, consistent with potent inhibition of CHIKV replication. In HEK293T cells (Figure 5B), RIBA maintained strong antiviral activity with minimal toxicity and Huh7 cells (Figure 5C) similarly displayed robust inhibition of viral replication by RIBA with minimal impact on confluence. Infection of all three cell lines generated substantial separation between mock and infected controls, with mean reporter signals approximately 37,000-fold, 20,500-fold and 4,900-fold higher in infected BHK-21, HEK293T and Huh7 cultures, respectively, demonstrating a large assay dynamic range and clear discrimination between infected and uninfected populations.

Across all cell types, these data demonstrate that the nsP3-tagged infectious clone platform enables simultaneous quantification of viral replication and host cell viability, facilitating discrimination between antiviral efficacy and compound-associated toxicity. In addition, the parallel development of the nsP3-mS3 infectious clone provides an orthogonal readout that could be deployed in cases where compound-associated green autofluorescence interferes with mNeonGreen-based detection, enhancing assay robustness and flexibility. Together, these findings demonstrate that the platform supports high-content, multi-parameter antiviral profiling across diverse cellular backgrounds, providing a foundation for evaluating how antiviral responses vary between cell types.

### 2.6. Primary human cells support nsP3-mNG CHIKV replication, but lose ribavirin antiviral phenotype

Primary HSkMMs and pHDFs have previously been identified as physiologically relevant models for CHIKV replication (Lentscher et al. 2020; Young et al. 2019; Lohachanakul et al. 2015; Hussain et al. 2016; Sourisseau et al. 2007; Taylor, Roberts, and Rayner 2026). We therefore assessed permissivity to nsP3-mNG CHIKV replication. Infection of HSkMMs at MOI 1 resulted in detectable mNG signal at early time points, with widespread monolayer infection evident by 18 hpi (Supp. Figure 2A). Replication in pHDFs displayed slower kinetics, however widespread monolayer infection was observed 24 hpi at MOI 10 (Supp. Figure 2B). This confirmed that HSkMMs and pHDFs were permissive to fluorescently-tagged CHIKV infection and support productive viral replication.

To determine whether the nsP3-mNG infectious clone platform could be applied to antiviral screening in physiologically relevant primary cells, HSkMMs and pHDFs were infected at an MOI of 1 or 10, respectively, in the presence of RIBA under conditions optimised from kinetic analysis. Intriguingly, no antiviral activity was observed in HSkMM cells across the concentration range tested (Figure 6A). A loss of antiviral effect following RIBA treatment was similarly observed in pHDFs (Figure 6B). Plaque assay analysis of supernatants harvested from HSkMMs at 18 hpi showed a comparable trend, with no change in viral titre following 20 µM RIBA treatment relative to untreated controls (Supp. Figure 3A).

**Figure 6:**
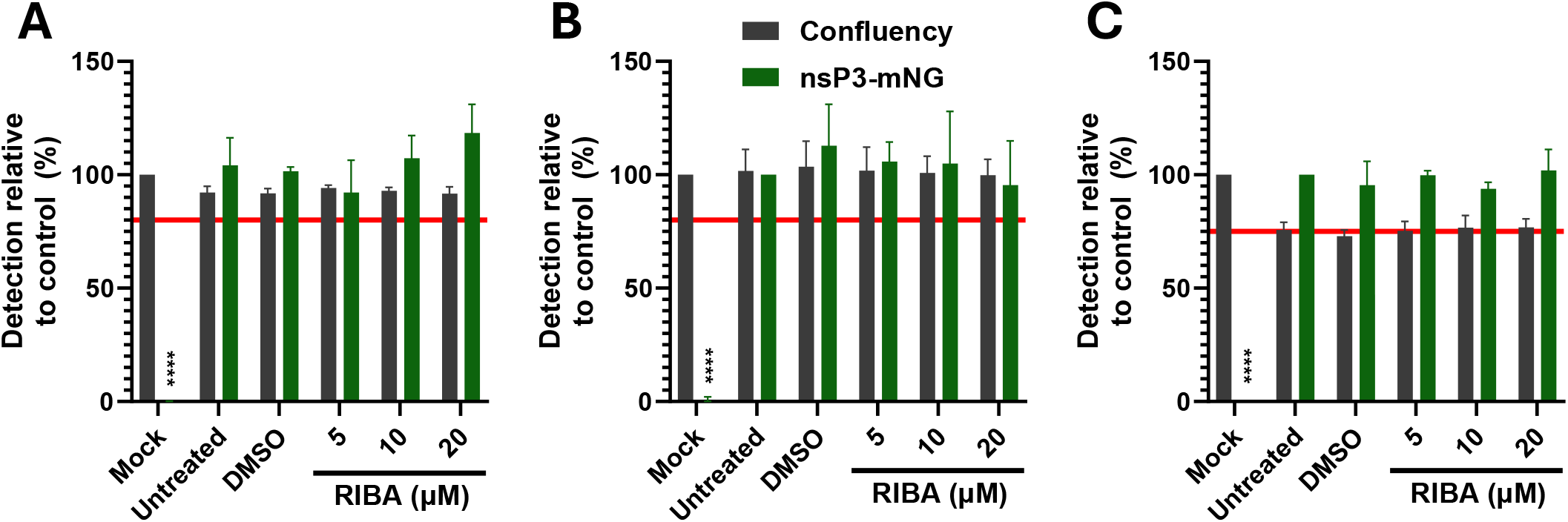
Assessment of antiviral activity and cytotoxicity in human skeletal muscle myoblasts (HSkMMs), primary- and hTERT-human dermal fibroblasts (pHDFs, hTERT-HDFs) using the CHIKV nsP3-mNG system. (A) HSkMMs were infected with CHIKV nsP3-mNG at an MOI of 1 in the presence of RIBA at 5, 10, and 20 µM prior to imaging at 18 hpi to assess viral replication. pHDFs (B) and hTERT-HDFs (C) were infected with CHIKV nsP3-mNG at an MOI of 10 in the presence of RIBA at 5, 10, and 20 µM prior to imaging at 24 hpi to assess viral replication. Cell confluency relative to mock control was evaluated as a measure of compound toxicity and mNG fluorescence was used as a measure of viral replication. Black asterisks denote statistically significant differences in mNG signal relative to untreated controls. Red line indicates the 80% relative confluency threshold for acceptable cell viability for HSkMM and pHDFs; in hTERT-HDFs the red line indicates the confluence of untreated, infected cells. Error bars for HSkMMs represent mean data ± SEM from two independent donors, each assessed in two independent experiments (biological n = 2, total n = 4) and n = 3 independent repeats for pHDFs and hTERT-HDFs.

The consistency between fluorescence-based measurements and plaque assay data indicates that mNG signal provides a reliable proxy for viral output in this system. Importantly, these results demonstrate that antiviral phenotypes observed in conventional screening cell lines do not necessarily translate to physiologically relevant primary human target cells. As such, the nsP3-mNG platform provides a valuable means of identifying context-dependent differences in antiviral activity and highlights the importance of physiologically relevant models for evaluating candidate inhibitors.

To bridge the gap between immortalised cell lines and primary HSkMMs, we next evaluated the nsP3-mNG CHIKV platform in hTERT-HDFs. These cells retain key features of primary human cells, including more intact innate immune signalling pathways, and as such provide an intermediate model for assessing antiviral response in a more physiologically relevant context than other transformed cell lines. hTERT-HDFs were first confirmed to support CHIKV replication (Supp. Figure 3C). At 8 hpi, diffuse cytoplasmic detection of nsP3-mNG was observed, with organisation into cytoplasmic foci later in infection (24 hpi) consistent with other cell lines.

To determine whether antiviral activity of RIBA observed in immortalised cell lines or primary HSkMMs and pHDFs were recapitulated in the intermediate cellular system, hTERT-HDFs were infected with nsP3-mNG, with read out 24 hpi (Figure 6C). Consistent with observations in HSkMMs and pHDFs, RIBA did not exhibit antiviral activity in hTERT-HDFs across the concentration range tested, with mNG fluorescence remaining comparable to untreated controls. This was further supported by plaque assay analysis, which showed no reduction in viral titres following treatment with 20 µM RIBA (Supp. Figure 3B).

### 2.7. Inhibition of nsP2 ablates replication in BHKs, HSkMMs, pHDFs and hTERT-HDFs

Given that RIBA had limited or no activity against CHIKV in HSkMMs, pHDFs and hTERT-HDFs, to validate the capability of the platforms to resolve mechanism-specific antiviral activity in the infectious virus systems, we evaluated the effects of RA.

In BHK cells, RA produced a dose-dependent reduction in mNG signal, with near-complete suppression observed at 1 and 10 µM (Figure 3E-H). No inhibition of viral replication was observed with 0.1 µM, consistent with SGR results at comparable time points (Figure 7A).

**Figure 7:**
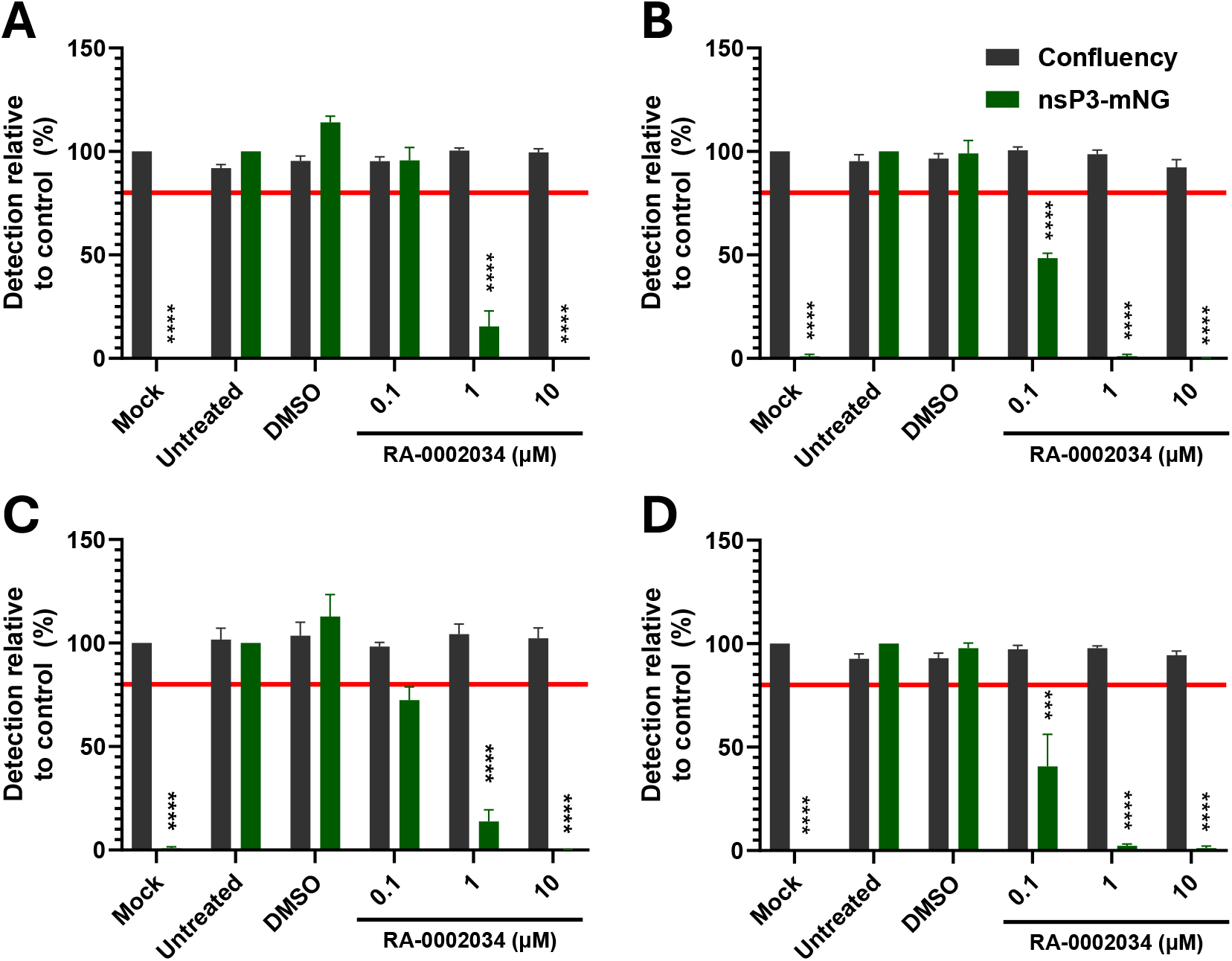
Assessment of antiviral activity of the nsP2 inhibitor RA-0002034 using infectious clone systems. BHK (A), HSkMM (B), pHDF (C) and hTERT-HDF (D) cells were infected with CHIKV nsP3-mNG (BHK, MOI 0.1; HSkMM, MOI 1; pHDF and hTERT-HDF, MOI 10) in the presence of RA at 0.1, 1 and 10 µM, and incubated for 16 hours (BHK), 18 hours (HSkMM) or 24 hours (HDF) prior to analysis. Cell confluency relative to mock control was evaluated as a measure of compound toxicity and mNG fluorescence was used as a measure of viral replication. Black asterisks indicate statistically significant differences in viral replication relative to untreated infected controls. Red line denotes the 80% confluency threshold for acceptable cell viability. Data are presented as mean ± SEM (n = 3 independent experiments).

In sharp contrast to RIBA, a markedly stronger response was observed in more physiologically relevant cell systems. In HSkMMs, RA treatment resulted in strong, dose-dependent suppression of viral replication, with near complete suppression at the higher concentrations of 1 and 10 µM (Figure 7B). pHDFs displayed a comparable phenotype at 1 and 10 µM, whilst 0.1 µM resulted in a more modest 28% inhibition. Comparable with other cell lines, hTERT-HDFs produced a substantial reduction in viral signal (Figure 7D). Across all cell types, antiviral effects were observed in the absence of reduction in cell confluency, indicating that reductions in fluorescence were not attributable to cytotoxicity.

These data demonstrate that RA acts as an effective inhibitor of CHIKV replication across all systems, consistent with targeting of nsP2-dependent replicase activity. Importantly, the stronger antiviral activity observed in primary and intermediate cell models relative to BHK-21 cells highlights the influence of cellular context on antiviral efficacy. Collectively, these data alongside that for RIBA, reinforce the observation that antiviral phenotypes identified in transformed cell lines are not necessarily retained in more physiologically relevant models under comparable experimental conditions. This highlights the capacity of the nsP3-mNG platform to resolve cell-type-dependent variation in antiviral activity, providing a framework for more predictive evaluation of candidate inhibitors.

### 2.8. Inosine supplementation and ENT1 inhibition do not restore CHIKV replication during RIBA treatment

To determine whether the mechanism of RIBA-mediated inhibition of CHIKV replication was linked to ENT1-associated nucleoside transport, BHK-21 cells were treated with RIBA in the presence of increasing concentrations of inosine. Inosine alone produced a dose-dependent reduction in viral replication, with the greatest effect observed at 1000 μM (Figure 8A). However, co-addition of inosine and RIBA failed to restore viral replication relative to RIBA-treated controls, indicating that inosine supplementation was unable to overcome RIBA-mediated inhibition.

**Figure 8:**
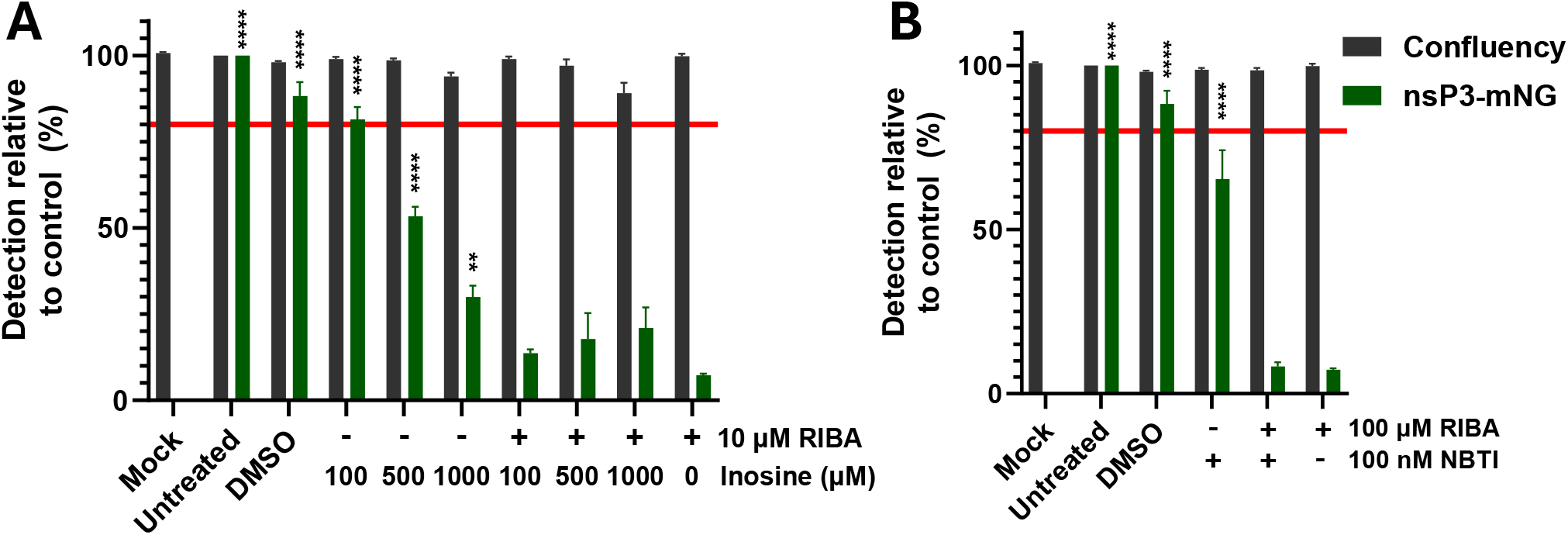
Inosine supplementation and ENT1 inhibition do not reverse ribavirin-mediated suppression of CHIKV replication. (A) BHK-21 cells were infected with CHIKV nsP3-mNG (MOI 0.1) and treated with 10 μM RIBA in the presence or absence of inosine (100, 500 or 1000 μM) before incubation for 16 h prior to analysis. (B) BHK-21 cells were pre-treated with 100 nM NBTI for 30 mins prior to infection with CHIKV nsP3-mNG (MOI 0.1). NBTI was maintained throughout infection. Following a 30 min incubation with virus inoculum, cells were treated with 10 μM RIBA and/or 100 nM NBTI and incubated for 16 h prior to analysis. Cell confluency relative to mock control was evaluated as a measure of compound toxicity and mNG fluorescence was used as a measure of viral replication. Black asterisks indicate statistically significant differences in viral replication relative to untreated infected controls. Red line denotes the 80% confluency threshold for acceptable cell viability. Data are presented as mean ± SEM (n = 3 independent experiments).

To further investigate the contribution of ENT1 activity, cells were pre-treated with the ENT1 inhibitor nitrobenzylthioinosine (NBTI). NBTI alone reduced CHIKV replication relative to untreated controls, suggesting that nucleoside transport contributes to efficient viral replication in BHK-21 cells. However, NBTI did not alter the antiviral activity of RIBA, with replication levels remaining comparable to those observed following RIBA treatment alone (Figure 8B).

Collectively, these findings demonstrate that neither inosine supplementation nor pharmacological inhibition of ENT1 is sufficient to reverse RIBA-mediated inhibition of CHIKV replication, suggesting that ENT1-dependent nucleoside transport is not the primary determinant of RIBA antiviral activity under these conditions.

### 2.9. IMPDH-associated depletion of guanine nucleotide pools is consistent with RIBA-mediated inhibition of CHIKV replication in BHK-21 cells

To investigate whether inhibition of guanosine nucleotide biosynthesis contributed to the antiviral activity of RIBA, BHK-21 cells were infected with CHIKV nsP3-mNG and treated with 10 μM RIBA or 1 μM of the IMPDH inhibitor mycophenolic acid (MPA) in the presence or absence of 100 μM guanosine. Both RIBA and MPA significantly reduced nsP3-mNG fluorescence relative to untreated controls indicating marked suppression of viral replication (Figure 9A). In contrast, supplementation with guanosine restored replication in both treatment groups to levels comparable to untreated controls.

**Figure 9:**
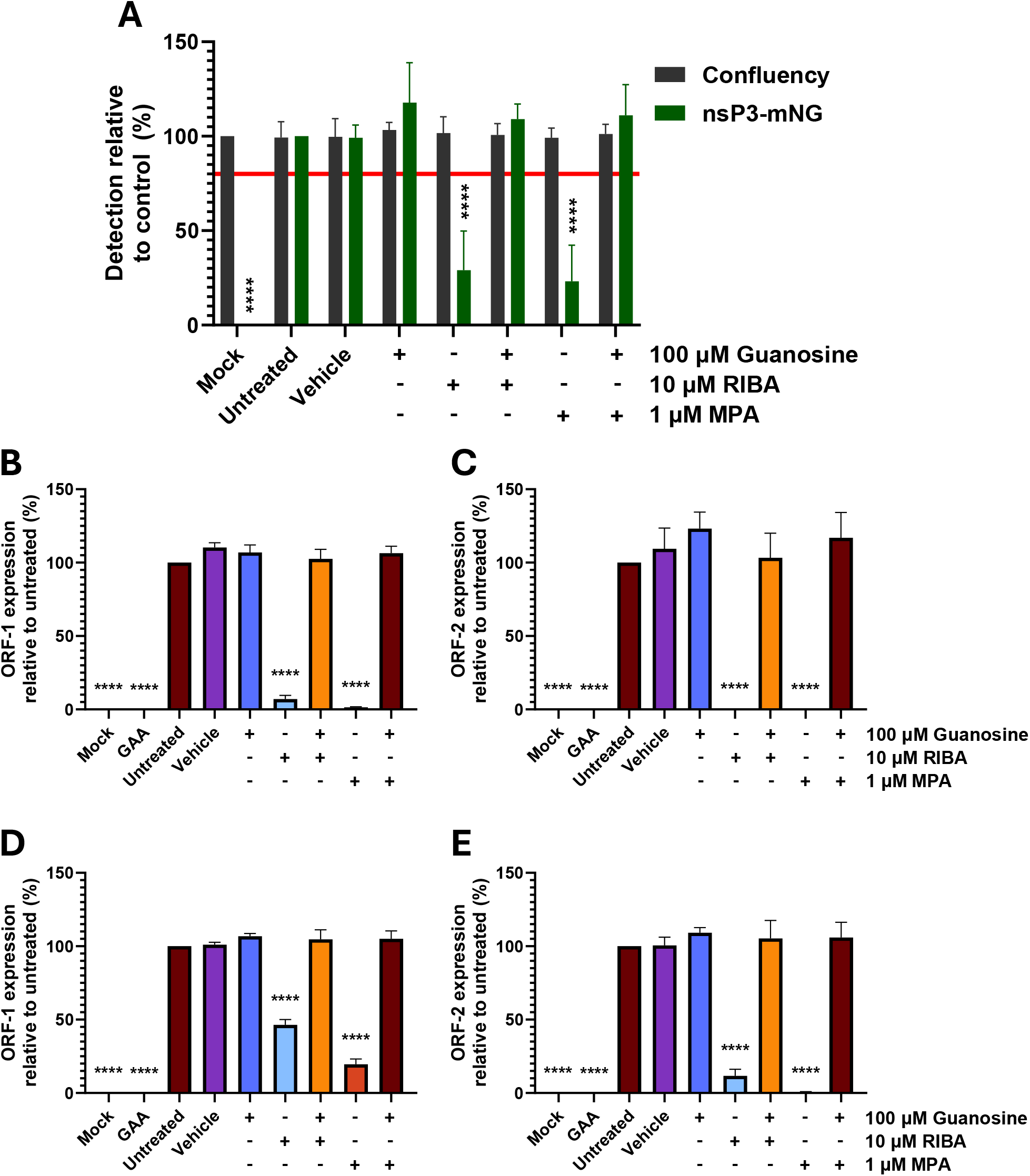
Ribavirin- and mycophenolic acid-mediated inhibition of CHIKV replication is rescued by guanosine supplementation in BHK-21 cells. (A) BHK-21 cells were infected with CHIKV nsP3-mNG (MOI 0.1) and treated with 10 μM RIBA or 1 μM MPA in the presence or absence of 100 μM guanosine and incubated for 16 h prior to analysis. Cell confluency relative to mock control was evaluated as a measure of compound toxicity and mNG fluorescence was used as a measure of viral replication. Black asterisks indicate statistically significant differences in viral replication relative to untreated infected controls. Red line denotes the 80% confluency threshold for acceptable cell viability. (B-E) BHK-21 cells were transfected with WT CHIKV sub-genomic replicon RNA in the presence of 10 μM RIBA or 1 μM MPA plus/minus 100 μM guanosine. DMSO was included as a vehicle control, alongside untreated, GAA and non-transfected (mock) controls. ORF-1 (nsP3-mNG) and ORF-2-mS3 expression were analysed at 8 hpt (B, C) and 16 hpt (D, E). Data are presented as mean ± SEM (n = 3 independent experiments).

To determine whether rescue occurred at the level of viral replication rather than during later stages of infection, the experiment was repeated using the dual-fluorescent CHIKV SGR. Consistent with the infectious virus assay, both RIBA and MPA reduced ORF-1 and ORF-2 expression at both 8 and 16 hpt (Figure 9B-E). Addition of guanosine restored reporter expression in all cases, reversing inhibition of both reporters and returning signal levels towards those observed in untreated controls. Rescue was observed at both early and late timepoints, indicating that replenishment of intracellular guanine nucleotide pools restored replication-associated processes throughout the course of replication.

Together with the inability of inosine supplementation to restore replication following RIBA treatment (Figure 8), these findings are consistent with IMPDH-associated depletion of intracellular guanine nucleotide pools as a major determinant of antiviral activity in BHK-21 cells. The observation that guanosine rescued both infectious virus replication and ORF-1- and ORF-2-associated reporter expression demonstrates that inhibition occurs upstream of structural gene expression and directly impacts the viral replication programme. More broadly, these data illustrate the ability of the dual-fluorescent platform to define the stage of the viral lifecycle affected by antiviral intervention, reinforcing its value as a mechanism-aware tool for antiviral discovery.

### 2.10. IMPDH inhibition exhibits limited antiviral activity in primary human cell models

To determine whether the antiviral phenotype associated with IMPDH inhibition was conserved in physiologically relevant human cell systems, HSkMMs and pHDFs were infected with CHIKV nsP3-mNG and treated with increasing concentrations of MPA (1-10 μM). Contrary to the pronounced inhibitory effects observed in BHK-21 cells (Figure 9), MPA exhibited minimal antiviral activity in primary cell models (Figure 10).

**Figure 10:**
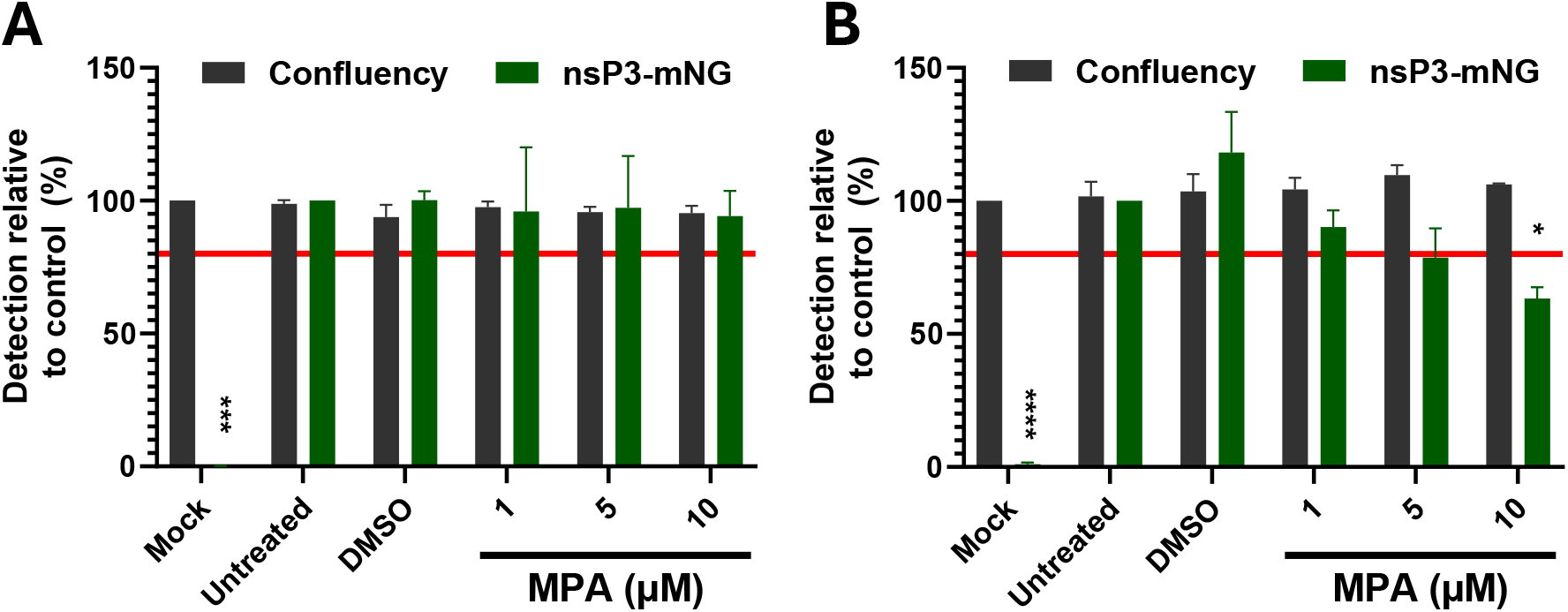
Mycophenolic acid does not substantially inhibit CHIKV replication in HSkMMs or pHDFs. HSkMMs (A) and pHDF (B) were infected with CHIKV nsP3-mNG (HSkMM, MOI 1; pHDF, MOI 10) and treated with 1, 5 and 10 μM MPA. Cells were incubated for 18 hours (HSkMMs) or 24 h (pHDFs) prior to analysis. Cell confluency relative to mock control was evaluated as a measure of compound toxicity and mNG fluorescence was used as a measure of viral replication. Black asterisks indicate statistically significant differences in viral replication relative to untreated infected controls. Red line denotes the 80% confluency threshold for acceptable cell viability. Data are presented as mean ± SEM (n = 3 independent experiments).

In HSkMMs, MPA treatment did not significantly reduce CHIKV replication at any concentration tested, with viral replication remaining comparable to untreated infected controls across the entire concentration range (Figure 10A). Similarly, pHDFs displayed only limited sensitivity to MPA, with no significant inhibition observed at 1 or 5 μM and only a modest 37% reduction at 10 μM (Figure 10B). Importantly, cell confluency remained above the predefined viability threshold under all conditions, indicating that observed effects were not attributable to compound-associated cytotoxicity.

These findings closely mirror the antiviral phenotypes previously observed for RIBA in primary cell systems (Figure 6), where potent inhibition detected in transformed cell lines was not retained in HSkMMs or fibroblast models. When considered alongside the guanosine rescue in BHK-21 cells (Figure 9), which demonstrated that both RIBA- and MPA-mediated inhibition were associated with depletion of intracellular guanine nucleotide pools, the reduced activity of MPA in primary cells suggests that susceptibility to guanine nucleotide depletion is highly dependent upon cellular context. Collectively, these data indicate that antiviral mechanisms readily identified in transformed screening systems may not be conserved in physiologically relevant human target cells and further emphasise the importance of incorporating primary cell models into CHIKV antiviral discovery pipelines.

## 3. Discussion

Understanding how interactions between CHIKV and its host cell regulate viral replication and antiviral susceptibility remains a major challenge in the field. Progress has been aided by the development of replicon and reporter-virus systems. However, existing approaches for studying CHIKV replication and evaluating antiviral activity are typically limited to endpoint-based readouts, such as luciferase-expressing replicons or immunofluorescence assays, which reduce viral replication to a single quantitative output and obscure underlying biological processes (Tsetsarkin et al. 2006; Colunga-Saucedo et al. 2022; Sawetpiyakul et al. 2025; Pohjala et al. 2011; Yin et al. 2026). In this study, we describe a dual-fluorescent SGR live-cell imaging platform that enables real-time, multiplexed interrogation of CHIKV replication dynamics. By coupling nsP3-mNG reporting of ORF-1 associated processes with mS3-based detection of sgRNA-dependent ORF-2 expression, this system simultaneously resolves distinct stages of the viral life cycle within a single assay. This platform was further extended into an infectious reporter virus system to permit complete lifecycle interrogation and overcome limitations inherent to replicon systems. Together, these complementary systems provide mechanistic resolution of CHIKV replication and enable direct assessment of how specific stages of the viral life cycle respond to genetic or pharmacological perturbation.

A key advance of the SGR platform is its ability to decouple genomic and sub-genomic translation events in real-time, providing mechanistic resolution not achievable using conventional single-reporter systems (Figure 1). Previous CHIKV replicon platforms have been used for antiviral screening but typically rely on luciferase or single fluorescent outputs that reflect aggregate replication activity. In contrast, the present system enables direct discrimination between replicase-associated processes and downstream structural gene expression. This was clearly demonstrated using the nsP3 PV247/8AA mutant, which selectively impaired sgRNA production while preserving ORF-1-associated replication activity, despite disrupted nsP3 scaffold formation (Figure 2) (Gao et al. 2019; Kril et al. 2024; Remenyi et al. 2018). The resulting divergence between nsP3-mNG and ORF-2-mS3 reporters confirmed that the system captured biologically distinct processes and provides a level of mechanistic resolution not accessible in existing CHIKV screening assays. Use of the PV247/8AA and GAA mutants further demonstrated the utility of the platform for assessing the effect of replication-associated mutations on viral fitness and reporter activity.

While SGR systems provide a powerful and biosafe approach for interrogating viral RNA translation and replication, they are inherently restricted to these stages of the CHIKV life cycle. Specifically, although replicons effectively model genome replication and sgRNA synthesis, they do not capture viral entry, assembly or egress. As such, compounds acting on these immediate-early or later stages cannot be identified using replicon-based systems alone, representing a key limitation of such approaches. To address this, we extended the platform to include fluorescently-tagged infectious CHIKV clones, allowing analysis to transition from replication-restricted systems to full life-cycle infection models (Figures 4, 6, 8). This provides an important bridge, linking mechanistic insight derived from the SGR system to biologically relevant outcomes during complete viral infection. The nsP3-mNG and nsP3-mS3 reporter viruses retained near-wild-type replication competence (Figure 4D-E), with only modest attenuation in viral titre and plaque morphology (Figure 4F-H), consistent with previous observations that the nsP3 hypervariable domain can accommodate insertions with limited disruption to replication (Remenyi et al. 2017; Pohjala et al. 2011).

Importantly, the infectious clone system enabled interrogation of processes not accessible in replicon models, including viral spread, reinfection and cytopathic progression. This distinction is particularly relevant for antiviral screening, where compounds that appear effective in replication-restricted systems may fail in the context of full viral infection. In addition, the availability of spectrally distinct reporters provides a practical advantage, by enabling mitigation of compound-associated autofluorescence and facilitating orthogonal validation of antiviral effects (Figure 4A-C). Together, the replicon and infectious clone systems establish a two-tier platform, spanning mechanistic dissection of RNA replication through to full viral infection within a unified imaging-compatible framework.

The utility of the SGR platform for antiviral screening was demonstrated using RIBA and RA, which exhibited dose-dependent inhibitory profiles in immortalised cell lines (Figure 3). RIBA produced strong inhibition of both ORF-1 and ORF-2 translation, while RA displayed partial inhibition at 0.1 μM, temporally restricted antiviral activity at 1 μM and complete suppression at 10 μM. These findings are consistent with previous reports. The ability of the platform to resolve these differences, further highlights the value of dual-reporter kinetic readouts for distinguishing mechanism-specific antiviral phenotypes.

Beyond development of the reporter systems, a key finding of this study was the pronounced cell-type dependence of antiviral activity (Figures 5-10). While strong antiviral effects were observed for RIBA in transformed cell lines, these effects were not retained in primary HSkMMs or hTERT-immortalised HDFs, where RIBA exhibited no detectable inhibition under optimised conditions (Figures 7, 8). This observation was supported by both fluorescence-based measurements and plaque assays, indicating that the lack of antiviral effect was not attributable to limitations of the reporter system. While previous studies reported anti-CHIKV activity for RIBA, such discrepancies are consistent with evidence that antiviral efficacy can vary substantially across host cell types (Franco et al. 2018; Shah, Sunderland, and Grdzelishvili 2010). Although the mechanistic basis for the reduced activity of ribavirin in primary human cells remains to be determined, differences in drug uptake, intracellular metabolism and innate immune signalling are known to influence antiviral activity, while CHIKV replication itself is strongly shaped by host-cell-specific interactions, particularly involving nsP3 and its associated replication complexes (Beaucourt and Vignuzzi 2014; Iikura et al. 2012).

Mechanistic interrogation of RIBA activity using the dual-colour platform provided additional biological insight. While RIBA has previously been reported to inhibit CHIKV replication, the dominant mechanism responsible for this activity remains incompletely resolved. Proposed explanations include inhibition of IMPDH and depletion of intracellular guanine nucleotide pools, direct effects on viral RNA synthesis and mutagenic activity following incorporation into nascent viral RNA. In the present study, supplementation with guanosine restored replication following treatment with both RIBA and the IMPDH inhibitor MPA, whereas inosine failed to rescue RIBA-mediated inhibition. Importantly, rescue was observed not only in infectious virus assays but also in the replication-restricted dual-colour SGR system, where restoration of both ORF-1- and ORF-2-associated reporter expression demonstrated recovery of replication-associated processes upstream of structural gene expression. Together, these findings are consistent with IMPDH-associated depletion of intracellular guanine nucleotide pools as a dominant determinant of RIBA activity under the conditions tested.

In contrast, evaluation of the nsP2 inhibitor RA demonstrated that physiologically relevant cell models remained highly permissive to antiviral intervention. RA exhibited stronger antiviral effects in HSkMMs and HDFs in comparison to BHK-21 cells, with substantial suppression of viral replication in both systems (Figure 9). This demonstrates that primary and intermediate cell models remain responsive to antiviral intervention and that the absence of RIBA activity does not reflect a general resistance to inhibition, but rather compound-specific variability in antiviral efficacy.

From a translational and antiviral discovery perspective, these findings are highly significant. While RIBA and MPA produced robust antiviral effects in transformed cell lines through a mechanism consistent with guanine nucleotide depletion, neither compound retained detectable activity in HSkMMs or dermal fibroblast models. This suggests that antiviral mechanisms identified using conventional screening systems may not be universally conserved across biologically relevant cellular models. Importantly, the ability of the platform to resolve both antiviral potency and underlying mechanism across multiple cellular contexts enabled identification of this discrepancy. These findings therefore highlight a critical challenge in antiviral discovery, namely that mechanistic vulnerabilities identified in transformed cell lines may not accurately predict the behaviour of candidate therapeutics in physiologically relevant target cells.

More broadly, the gradient of antiviral responses observed across transformed cells, HDFs and HSkMMs highlights the importance of incorporating multiple cellular systems into antiviral discovery pipelines. By integrating mechanistic resolution with comparative analysis across transformed, immortalised and primary cell systems, the platform enables simultaneous assessment of antiviral potency, mechanism-of-action and physiological relevance within a single experimental framework.

Collectively, these findings establish the platform as a high-content, mechanistically informative system for investigating CHIKV replication and antiviral responses, while demonstrating that antiviral responses can differ substantially between transformed and physiologically relevant human cell models. By integrating dual-reporter resolution, real-time imaging and applicability across a spectrum of cellular models, the system enables simultaneous assessment of antiviral activity and physiological relevance. Importantly, the platform identifies not only antiviral activity, but also the mechanisms underpinning that activity and their failure to translate into physiologically relevant cellular systems. The ability to connect antiviral efficacy with specific stages of the CHIKV replication programme represents a significant advance over existing endpoint-based screening approaches and provides a more predictive framework for investigating antiviral mechanisms in biologically relevant cell models.

Future applications of this system include large-scale compound screening, host-factor interrogation and detailed investigation of replication complex dynamics. Integration with automated image analysis and adaptation to higher-density formats in permissive systems may further enhance throughput. More broadly, the principles described here are readily transferable to other alphaviruses and viruses, providing a platform for enhancing antiviral discovery.

## 4. Materials and Methods

### 4.1. Cell culture and maintenance

Huh7 (human hepatocellular carcinoma), BHK-21 (baby hamster kidney), HEK293T (human embryonic kidney), primary adult human dermal fibroblasts (pHDFs, purchased from Caltag Medsystems) and hTERT-immortalised human dermal fibroblast (hTERT-HDF) cells were maintained in Dulbecco’s modified Eagle’s medium (DMEM, Sigma) containing 10% (v/v) foetal bovine serum (FBS, Sigma), 100 U/mL penicillin and 100 mg/mL streptomycin (Sigma). Huh7 medium was additionally supplemented with non-essential amino acids (Cytiva). For generation of hTERT-HDFs, adult dermal Fibroblasts were grown out from 3 mm punch biopsies and immortalised with hTERT as previously described (Gillespie et al. 2018).

Human Skeletal Muscle Myoblasts (HSkMM, Lonza, cat. no. CC-2580) were cultured in Sk-GM-2 growth medium (Lonza, cat. no. CC-3245) according to manufacturer’s instructions.

All cells were maintained at 37 °C in 5% CO_2_ and were routinely tested for mycoplasma contamination using a MycoAlert assay (Lonza).

### 4.2. Small molecule inhibitors

RIBA (cat no. R9644), inosine (cat no. I4125), guanosine (cat no. G6264), and mycophenolic acid powder (cat no. M3536) were purchased from Merck. RA-0002034 (cat no. 42186) and nitrobenzylthioinosine (cat no. T8551) were purchased from Cambridge Bioscience. All inhibitors were dissolved in DMSO stored at –20 °C until use, with the exceptions of inosine (dissolved in H_2_O prior to storage) and guanosine (dissolved in H_2_O fresh before use).

### 4.3. Plasmid construction

The CHIKV infectious clone rescue (ICRES) and SGR plasmids were derived from the CHIKV ECSA strain isolate LR2006 OPY1 (accession number DQ443544) (Tsetsarkin et al. 2006). WT and GAA CHIKV dual-fluorescent replicons were generated using NEBuilder HiFi DNA Assembly (NEB) with PCR products amplified from CHIKV dual luciferase plasmids spanning the tag insertion sites (Gao et al. 2019), 3xnls-mNeonGreen and pDx_mScarlet-I3. 3xnls-mNeonGreen (Addgene plasmid # 98875; http://n2t.net/addgene:98875; RRID:Addgene_98875) and pDx_mScarlet-I3 (Addgene plasmid # 189757; http://n2t.net/addgene:189757; RRID:Addgene_189757) were gifts from Dorus Gadella (Chertkova et al. 2020; Gadella et al. 2023).

The mScarlet-I3 amplicon was generated by overlap PCR to introduce a silent nucleotide substitution (c.510C>A), preserving a unique NotI restriction site for linearisation. The nsP3 PV247/8AA double amino acid substitution was inserted into the WT SGR by overlap PCR followed by restriction digest.

The nsP3-mNG ICRES was generated by transferring the tagged sequence into WT ICRES via restriction digest. The nsP3-mS3 ICRES was generated similarly from an unpublished nsP3-mS3 ORF-2-luc construct.

Primer sequences are available upon request.

### 4.4. *In vitro* transcription of viral plasmids

CHIKV SGR and ICRES plasmids were linearised using NotI-HF (NEB) and purified using the Monarch PCR & DNA cleanup kit (NEB). *In vitro* transcription was performed using the mMESSAGE mMACHINE SP6 transcription kit (Thermo Fisher Scientific) according to manufacturer’s instructions, within incubation at 37 °C for 2 h. Synthesised RNA was purified using the PureLink RNA mini kit (Thermo Fisher Scientific), with concentration determined by NanoDrop (Thermo Fisher Scientific) and integrity confirmed by MOPS-formaldehyde agarose gel electrophoresis. RNA was stored at −80 °C until use.

### 4.5. Sub-genomic replicon assays

BHK-21 cells were seeded at a density of 1 x 10^5^ cell/well in 24-well plates 16 h prior to transfection. 1 mg RNA and 3 mL Lipofectamine MessengerMAX reagent (Thermo Fisher Scientific) were separately diluted in 25 mL opti-MEM and incubated for 5 min at room temperature. Solutions were then combined and incubated for a further 15 min at room temperature. Medium was aspirated from cells, cells were washed once with PBS, and 450 mL phenol red-free DMEM supplemented with GlutaMAX and 10% FBS (complete DMEM, cDMEM) was added to each well. A total of 50 mL RNA:lipofectamine complex was added dropwise to wells and mixed gently. Live-cell imaging was performed hourly using an IncuCyte S3 system.

For screening assays, BHK-21 cells were seeded at a density of 1 x 10^4^ cells/well in 96-well plates. Small molecule inhibitors (SMIs), or equivalent volume of DMSO as a vehicle control, were diluted to 2X experimental concentration in 50 mL cDMEM in 96-well plates. For transfection, master mixes were prepared by diluting 150 ng RNA or 0.45 mL Lipofectamine MessengerMAX in 5 mL opti-MEM per well. The mixtures were incubated for 5 min at room temperature prior to combination, followed by a further 15 min incubation at room temperature to permit complex formation. 40 mL cDMEM was added per well and mixed by pipetting. 50 mL transfection mix was added to each well to dilute compounds to 1X, after which cells were washed once with PBS and 100 mL transfection/SMI mixture was added to each well. Hourly live-cell imaging was performed using an IncuCyte S3 system.

### 4.6. Virus propagation

Propagation of virus stocks from plasmid was performed as described previously using BHK-21 cells (Sun et al. 2024). Once cytopathic effect (CPE) was observed (30-48 h post-infection, hpi), supernatant was collected and clarified by centrifugation at 2,000 *g* for 5 min to remove debris, and viral titre quantified by plaque assay.

### 4.7. Viral quantification by plaque assay

BHK-21 cells were seeded at a density of 2.5 x 10^5^ cells/well in a 12-well plate. Medium was removed and cell monolayers were infected in duplicate as previously described with 10-fold serial dilutions of CHIKV (10^-3^ to 10^-8^). Following binding, inoculum was removed and replaced with a 0.8% methylcellulose DMEM overlay for 72 h. Monolayers were fixed with 4% (v/v) formaldehyde PBS solution prior to staining with crystal violet. Plates were imaged using a GelDoc Go System (Bio-Rad), and plaques were counted to calculate viral titre as PFU/mL (plaque-forming units per mL). Plaque size was determined using the ViralPlaque ImageJ plugin (Cacciabue, Curra, and Gismondi 2019).

### 4.8. CHIKV infection assays

BHK-21 (1 x 10^5^) and HDF (5 x 10^4^) cells were seeded 16 h prior to infection in 24-well plates. HSkMM cells were seeded at a density of 1.8 x 10^4^ cells/cm^2^ in 24-well plate 48 h prior to infection. Unless otherwise stated, virus stocks were diluted to desired MOI (BHK-21, 0.1; HDF, 10; HSkMM, 1) in cDMEM (BHK-21), cDMEM containing 2% FBS (HDF) or phenol-free DMEM/F12 supplemented with GlutaMAX and 2% FBS (HSkMM). Growth medium was aspirated from wells, and virus-containing medium added. Plates were rocked at 15 rpm for 5 min, followed by incubation at 37 °C for 30 mins to facilitate virus adsorption. Infection medium was then replaced with fresh prewarmed medium, and cells were incubated for desired time. For nsP3-mNG and nsP3-mS3 tagged viruses, hourly live-cell imaging was performed using an IncuCyte S3 instrument at 10X magnification. For plaque assays, supernatants were harvested 16 hpi (BHK-21) or 24 hpi (HSkMM and HDF) and clarified by centrifugation at 2,000 *g* for 5 mins to remove cellular debris prior to titration.

For blind passaging experiments, supernatants were harvested 18 hpi and clarified by centrifugation at 2,000 *g* for 5 mins. Naïve monolayers were infected with 1 μL viral supernatant diluted in fresh growth medium as described above, with analysis of fluorescence at 18 hpi.

For screening assays, BHK-21 (1 x 10^4^), Huh7 (1 x 10^4^) and HEK293T (1.5 x 10^4^) cells/well were seeded in 96-well plates, and 5 x 10^4^ HDF cells/well were seeded in 24-well plates 16 h prior to infection. HSkMM cells were seeded at a density of 1.8 x 10^4^ cells/cm^2^ in 24-well plate 48 h prior to infection. SMIs were diluted to 2X experimental concentration in 50 mL (96-well assay) or 250 mL (24-well assay). An equivalent volume of diluted nsP3-mNG virus stock was added to achieve the final MOI (BHK-21 and HEK293T, 0.1; HSkMM, 1; Huh7 and HDF, 10). Growth medium was aspirated, and the inhibitor-virus mixture was added to the cells. Plates were rocked at 15 rpm for 5 min, followed by incubation at 37 °C, with hourly live-cell imaging performed using an IncuCyte S3 system at 10X magnification.

### 4.9. Western blotting

For western blotting, BHK-21 cells were lysed at 16 hpi using 1X Laemmli buffer and DNA sheared by passing through a 21-gauge needle. Following SDS-PAGE, samples were transferred onto a PVDF membrane using a Trans-Blot Turbo System (Bio-Rad) at 25 V for 40 mins. Blocking and antibody incubations were performed using 5% skimmed milk (w/v) in TBS-T. Three TBS-T washes were performed after primary and secondary antibody incubations.

Antibodies targeting nsP3 (1:1,000, kind gift from Andres Merits), mNeonGreen (1:1,000, ProteinTech, cat. no. 32F6), mCherry (1:2,000, ProteinTech, cat. no. 26765-1-AP) or GAPDH (1:1,000, Santa Cruz Biotechnology, cat. no.sc-32233) were diluted in 5% (w/v) skimmed milk in TBS-T. Membranes were incubated with diluted antibody solution overnight at 4 °C with agitation. Anti-mouse 680 (cat. no. 35518) and anti-rabbit 800 (cat. no. SA5-35571) DyLight secondary antibodies were purchased from Thermo Fisher Scientific, diluted 1:5,000 in 5% (w/v) skimmed milk in TBS-T and incubated with membranes for 1 h at room temperature with agitation protected from light. Blots were air-dried, protected from light, before detection using a Li-Cor Odyssey M Imaging System.

### 4.10. IncuCyte analysis

Image analysis of nsP3-mNG and nsP3-mS3 infection assays was performed using the IncuCyte 2025B Basic Analyzer software. Phase confluence was determined using AI confluence segmentation. Green fluorescence was quantified using surface fitting segmentation, edge splitting enabled, sensitivity −35, green calibrated unit threshold 2.0 and area minimum 25 mm^2^. Red fluorescence was analysed using equivalent parameters, with a red calibrated threshold 0.5.

For SGR assays green parameters were unchanged, whereas red fluorescence was quantified using surface fitting segmentation with edge splitting enabled, sensitivity 0, red calibrated unit threshold 10.0 and area minimum 50 mm^2^.

### 4.11. Statistical analysis

Statistical analysis was performed using GraphPad Prism 10 with one-way ANOVA unless otherwise stated. *P* value £0.05=*, <0.01=**, <0.001=*** and <0.0001=****.

## 5. Author contributions

SJD and AKT conceived and designed the study. SJD and MGAB performed the experiments. SJD, MGAB and LW analysed the data under supervision of AKT. FDG and CWW provided essential resources. Funding was acquired by AKT. SJD wrote the original draft of the manuscript. All authors reviewed, edited and approved the final manuscript.

## 7. Data access statement

The datasets generated and/or analysed during the current study are available from the corresponding author upon reasonable request.

## 10. Acknowledgements

The authors gratefully acknowledge Professor Andres Merits (University of Tartu) for his foundational contributions to the CHIKV sub-genomic replicon and infectious clone systems that enabled this study. We further thank him for generously providing key reagents and for his insightful discussions and critical review of the manuscript.

---

**Supplementary Figure 1:**
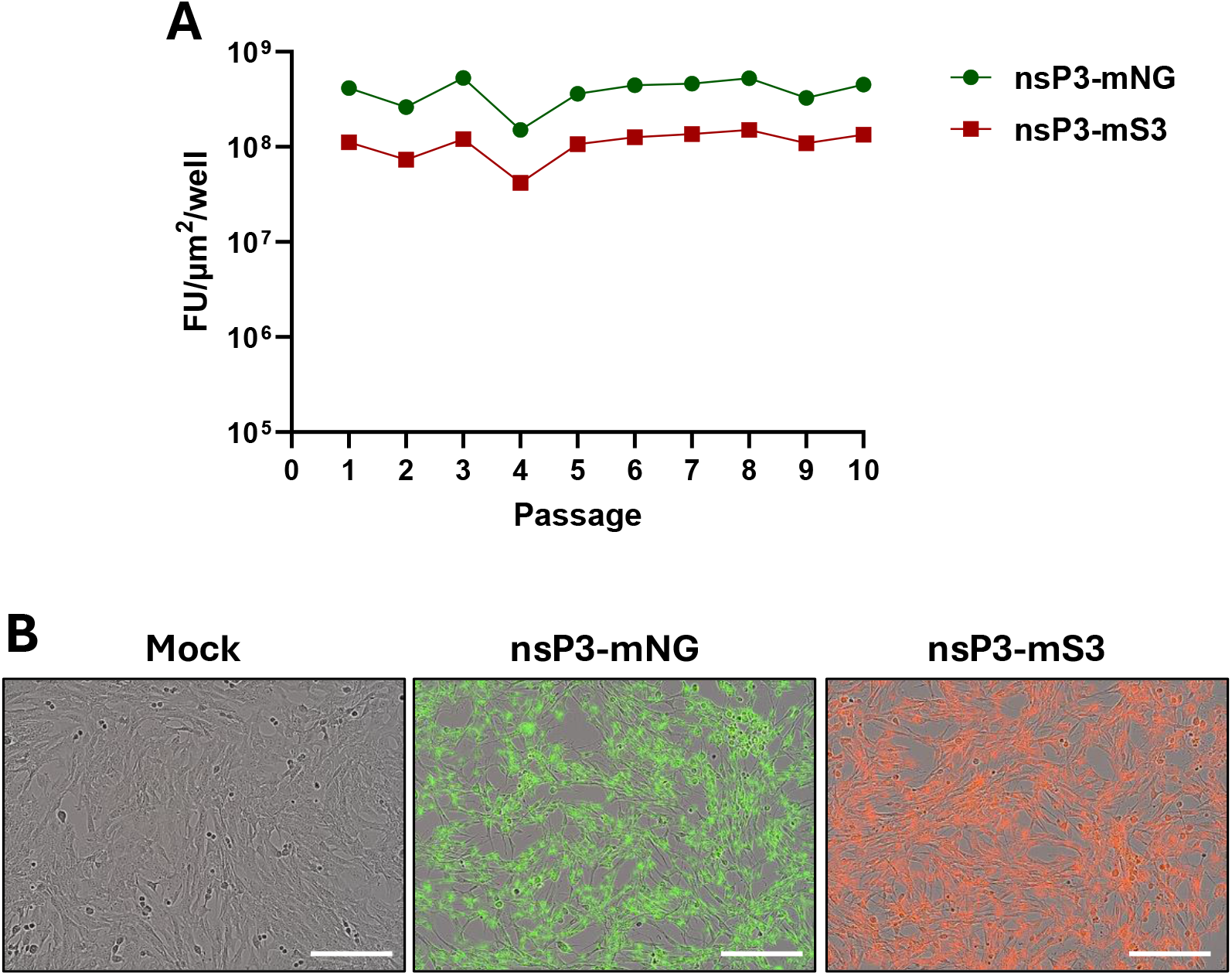
nsP3-tagged infectious clones retain fluorescence across multiple passages. (A) Quantification of nsP3-mNG and nsP3-mS3 reporter fluorescence following blind passage in BHK-21 cells. Fluorescence was analysed by live-cell imaging 18 hpi and is quantified as fluorescence units/µm^2^/well (FU/µm^2^/well). (B) Infection of BHK-21 cells with nsP3 reporters to confirm complete monolayer infection. Scale bars 400 µm.

**Supplementary Figure 2:**
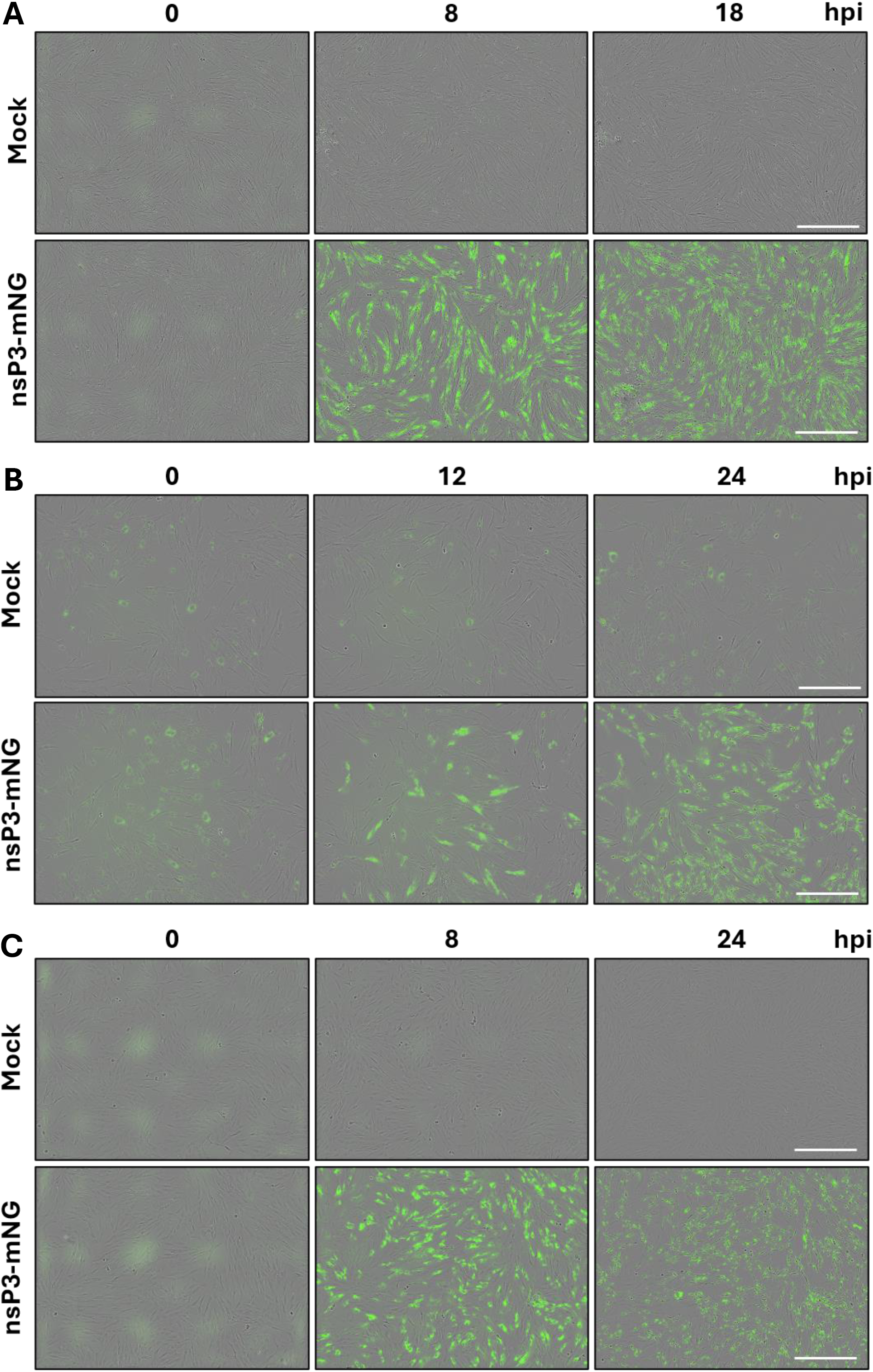
Primary HSkMM, pHDFs, and hTERT-immortalised HDFs are permissive to CHIKV nsP3-mNG infection. HSkMM (A), pHDF (B) and hTERT-HDFs (C) were infected with CHIKV nsP3-mNG and monitored by live-cell imaging. Representative phase-contrast/fluorescence overlay images acquired at increasing time points post infection. Scale bars 400 µm.

**Supplementary Figure 3:**
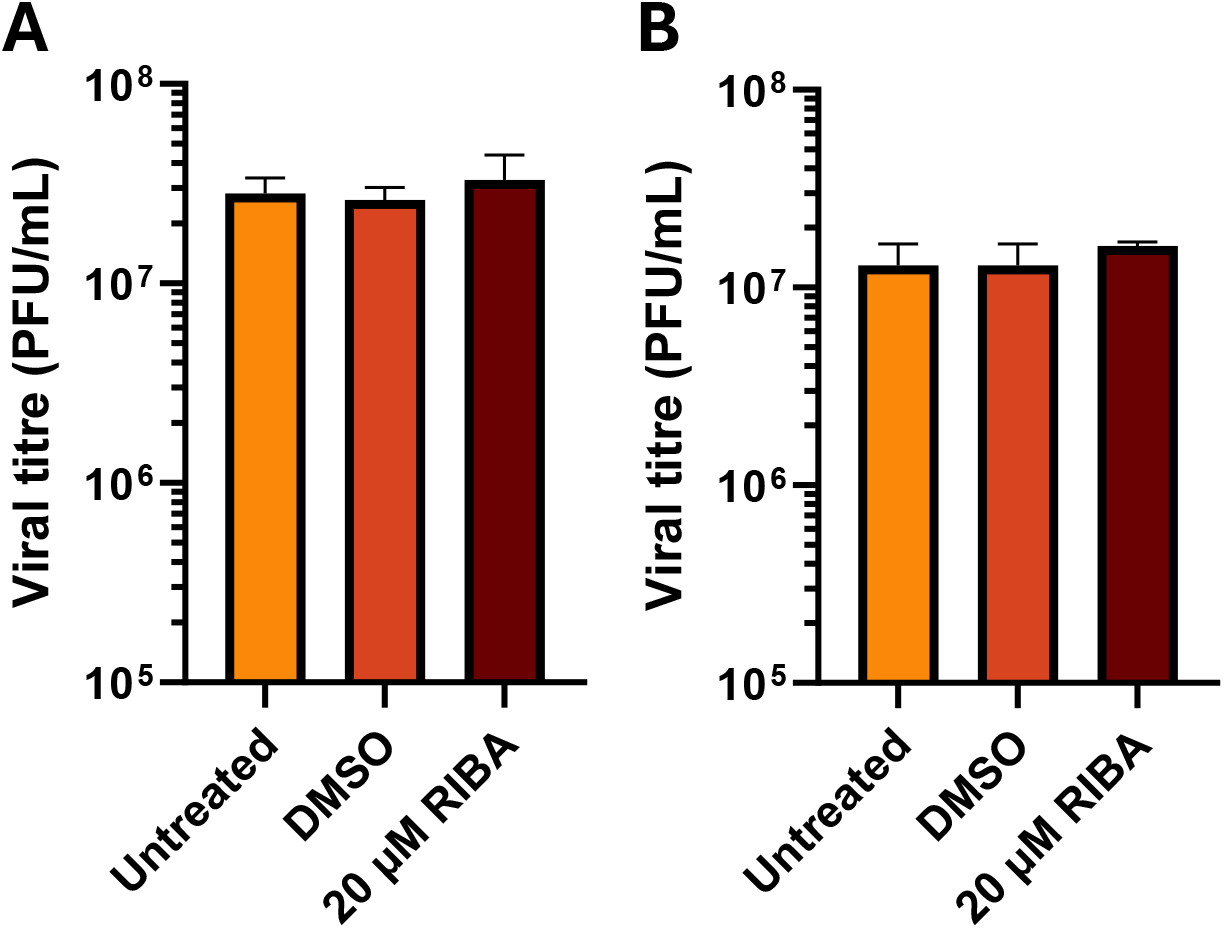
Assessment of ribavirin antiviral activity in HSkMMs and hTERT-HDFs by plaque assay. Supernatants harvested 18 hpi (HSkMMs) and 24 hpi (pHDFs) were evaluated by plaque assay to determine viral titre in HSkMMs (A) and hTERT-HDFs. No statistical significance was observed between samples. Data are presented as mean ± SEM (n = 3 independent experiments).

